# Poxvirus A52 is a host range factor for modified vaccinia virus Ankara (MVA) and promotes viral replication by disturbing the formation of autolysosomes

**DOI:** 10.1101/2024.06.13.598619

**Authors:** Kang Niu, Yongxiang Fang, Yining Deng, Ziyue Wang, Shijie Xie, Junda Zhu, Baifen Song, Wenxue Wu, Zhizhong Jing, Chen Peng

## Abstract

Many members of the poxvirus family are important zoonotic pathogens that pose a significant threat to human and animal health worldwide. Autophagy is a multi-step degradation pathway within cells, and one of its primary biological functions includes the clearance of invading viruses. Nevertheless, the interplay between poxviruses and host cell autophagy has not been fully elucidated. Here, we demonstrate that vaccinia virus (VACV) and lumpy skin disease virus (LSDV) induce incomplete autophagy and inhibit the fusion of autophagosomes and lysosomes, while modified vaccinia virus Ankara (MVA), an attenuated strain of VACV unable to replicate in almost all human cells, does not. Additionally, we screened and identified the VACV protein A52 as a key factor that obstruct the formation of autolysosomes. Mechanistically, A52 interacts with SNAP29 and inhibits its interaction with STX17 and VAMP8, both of which are binding partners of SNAP29 and are essential for complete autophagy. Moreover, A52 promotes the proteasomal degradation of SNAP29, which facilitates viral replication. We further revealed that SNAP29 functions as a restriction factor for MVA, as the suppression of SNAP29 allowed the replication of MVA in human cells. In summary, our data present a molecular mechanism by which poxviruses manipulate the cellular autophagic machinery and provide additional explanation for the restriction of MVA in human cells.

## 1. INTRODUCTION

Poxviruses, including mpox, lumpy skin disease virus (LSDV) and sheeppox virus, continue to pose a significant threat to human and animal health after the successful eradication of smallpox through extensive vaccination efforts^1, 2^. Mpox has resulted in infections in over 92,783 individuals across more than 100 countries^3^. LSDV, a member of the *Capripoxvirus* genus, leads to significant economic losses in the cattle industry in affected countries due to decreased milk production and increased mortality rates resulting from its infection^4^. Vaccinia virus (VACV) is the most well-studied prototype poxvirus and the vaccine strain used to eliminate smallpox. Investigating how poxviruses exploit host cell machinery and employ viral strategies to evade host immune responses is crucial for the development of future vaccines and therapeutic interventions. Modified vaccinia virus Ankara (MVA), an FDA-approved vaccine strain for smallpox and mpox, was generated through repeated passages of its parental strain chorioallantois vaccinia virus Ankara (CVA) in chicken embryo fibroblasts (CEF)^5^. MVA has lost its ability to replicate in most mammalian cells, including almost all human cells^6^. The underlying mechanism is complex and has only been partially elucidated by recent studies^7–10^. However, the mechanisms that restrict its replication in human cells remain intricate and require further investigation.

Autophagy, an evolutionarily conserved, multi-step intracellular protein degradation process^11, 12^, is involved in a wide array of physiological and pathological processes, including the defense against viral infections^13–18^. Nevertheless, its exact role in controlling viral infections remains ambiguous, as both beneficial and detrimental effects have been observed across different types of viruses. Macroautophagy includes three critical steps: 1) initiation of nucleation; 2) extension of phagocytic vesicle structure to seal the cytoplasmic components and the formation of complete autophagosomes; 3) the fusion of autophagosomes with lysosomes to form autolysosomes, which then leads to degradation of encapsulated contents^19–21^. All these steps are tightly regulated and reports have shown various strategies employed by viral pathogens to hijack or prohibit all or partial of these processes^18^. Emerging evidence has shown that many viruses have evolved mechanisms to evade or inhibit one or multiple steps of the autophagic pathway for their own benefit^22–24^. For example, the interaction between the human simplex virus 1 (HSV-1) ICP34.5 and cellular Beclin 1 facilitates the inhibition of autophagy and increases viral replication^25^; the human cytomegalovirus (HCMV) encodes a homolog of ICP34.5, namely TRS1, which also inhibits autophagy through beclin 1 interaction to promote viral replication^26^; the K7 protein of Kaposi’s sarcoma virus (KSHV) promotes Rubicon-Beclin 1 interaction, which results in the inhibition of VPS34’s enzyme activity and enhances viral survival^27^; the vFLIP protein of KSHV inhibits autophagy by blocking the binding of ATG3 and LC3 during autophagosome elongation^28^.

On the other hand, viruses can exploit the autophagic machinery for their replication. The double membrane compartment formed during autophagy can provide a physical platform for viruses to replicate by concentrating viral intermediate products, and protection of viral components from immune surveillance and degradation. E.g., treatment of cells with an autophagy inducer rapamycin (rapa.) increased the replication of poliovirus, while silencing of autophagic factors reduced viral replication^29^. In addition, electron microscopic analysis of poliovirus-infected cells showed DMVs (double-membrane vesicles) resembling autophagosomes provided a scaffold for viral RNA replication. Previous reports showed deletion of several autophagy factors (ATG3, ATG5, Beclin 1) failed to dampen VACV’s replication in mouse embryonic fibroblast (MEF) cells ^30, 31^. However, the exact role of autophagy during poxvirus infection in human cells and strategies employed by poxviruses to manipulate autophagy, remain enigmatic.

Through a screening for small-molecule antivirals, we identified several autophagic inhibitors that exhibited strong inhibitory capability on VACV replication. Further investigation suggested manipulation of autophagy might affect VACV’s replication as pharmaceutical inhibition and induction of autophagy led to reduced and increased replication of several poxviruses, respectively. Interestingly, although autophagy activation was beneficial for VACV and LSDV’s replication, both viruses were able to inhibit the final step of autophagy, namely the formation of autolysosomes, while MVA failed to do so. We further identified that VACV protein A52 was responsible for the inhibition of autolysosome formation. Our data further characterized the mode of action by which A52 blocked autolysosomes formation and identified its cellular target SNAP29. As the insertion of A52 into MVA partially improved its replication in human cells, A52 is thus considered a novel host range factor for MVA. Moreover, ectopic expression of SNAP29 inhibited the replication of MVA or a strain of VACV that lacks A52R, suggesting the host restriction role of SNAP29 for MVA. Our findings demonstrate the importance of autophagy in promoting poxvirus replication, report a novel restriction pathway for MVA in human cells, and characterize the detailed mechanism by which VACV blocks the fusion of lysosomes with autophagosomes to promote its replication.

## 2. RESULTS

### 2.1 Autophagic inhibition suppresses the replication of VACV

To determine whether inhibition or activation of autophagy modulates VACV replication, A549 or DF1 cells were infected with VACV-Western Reserve (WR) or MVA at 3 PFU/cell, respectively, in the presence or absence of various dosages of autophagy agonist rapamycin (rapa.), or antagonist bafilomycin A1 (baf. A1). Viruses were harvested 24 hours post infection (hpi) for titration. The results showed autophagy agonist rapa. promoted the replication of VACV-WR and MVA in a dose-dependent manner, while baf. A1 effectively suppressed viral replications (Figures 1A and 1B). The chemicals showed a similar effect on the replication of LSDV (Figures S1A and S1B). To explore if pharmaceutical inhibition or activation of autophagy affected viral mRNA abundance and DNA replication, A549 cells were infected with VACV-WR in the presence of rapa. or baf. A1 and total RNA and DNA were harvested for mRNA and viral genomic DNA quantification using quantitative realtime-PCR (qRT-PCR)^32^. Cytosine arabinoside (AraC) was included as a positive control as it was able to ablate viral post-replicative gene expression^33^. To our surprise, mRNA levels of selected viral early (E3L), intermediate (D13L) and late genes (A3L), as well as viral DNA replication were not significantly affected by rapa. or baf. A1 treatment, although AraC abolished both viral DNA replication and post-replicative gene transcription (Figures 1C-1F). Next, we investigated the effect of autophagic induction or inhibition on viral protein synthesis. Human A549 cells were treated with rapa., Earle’s Balanced Salt Solution (EBSS), or baf. A1 prior to VACV-WR infection at 3 PFU/cell and total protein was collected at 12 hpi for Western blotting analysis and viral quantification. Although the synthesis of viral I3, an early protein, was not affected by autophagic induction, the synthesis of A3 was slightly enhanced by the treatment of EBSS and rapa., both of which are autophagy inducers (Figure 1G). Similarly, the synthesis of LSDV-H3 (a late protein) was enhanced by the treatment of EBSS and rapa. (Figure S1C). And the use of these chemicals slightly enhanced viral replication (Figures H and S1D). In contrast, the addition of autophagic inhibitor baf. A1 evidently suppressed A3 synthesis and viral replication (Figures 1I and 1J). Similarly, the addition of baf. A1 suppressed LSDV-H3 synthesis and viral replication (Figures S1E and S1F). These results suggested inhibition or activation of autophagy inhibited or enhanced viral late protein synthesis and viral replication. As small-molecule chemicals may present off-target effects, we next verified our observation by suppressing various factors important for autophagy regulation, including ATG3, ATG7, ATG12, Beclin 1 and ATG16L1^34^. Human A549 cells were transfected with siRNA targeting the above-mentioned genes as well as si-NC as a negative control for 48 h prior to VACV-WR infection and viral progenies were collected at 24 hpi and quantified by the method described above. As shown in Figure S1G, Figure S1I and Figure S1K, transfection of siATG3, siATG7 and siATG12 effectively reduced the levels of corresponding ATG proteins, but failed to influence viral titers (Figures S1H, S1J and S1L). Nevertheless, suppression of ATG16L1 and Beclin 1 significantly inhibited viral A3 synthesis (Figures 1K and 1M) as well as viral replication (Figures 1L and 1N). Similarly, suppression of ATG16L1 and Beclin 1 inhibited LSDV-H3 synthesis (Figures S1M and S1O) as well as viral replication (Figures S1N and S1P). Next, we generated a cell line stably expressing shATG16L1, as well as its revertant control expressing a KD-resistant ATG16L1 and further examined viral protein synthesis and viral replication within these cells. In agreement with our previous observation, viral A3 expression was reduced (Figure 1O) and the titers of VACV-WR were reduced by approximately 10× folds (Figure 1P) when ATG16L1 was stably knocked down compared to that in the control cells, but the reduction was compensated in the revertant control, in which the expression of ATG16L1 was rescued (Figures 1O and 1P). Similar results were also observed for LSDV (Figures S1Q and S1R). In summary, these data demonstrated that inhibition of autophagy by baf. A1 or suppression of ATG16L1 and Beclin 1 in cells was able to prohibit replication of VACV, MVA and LSDV, whereas the pharmaceutical induction of autophagy promoted viral replication within cells.

**Figure 1.**
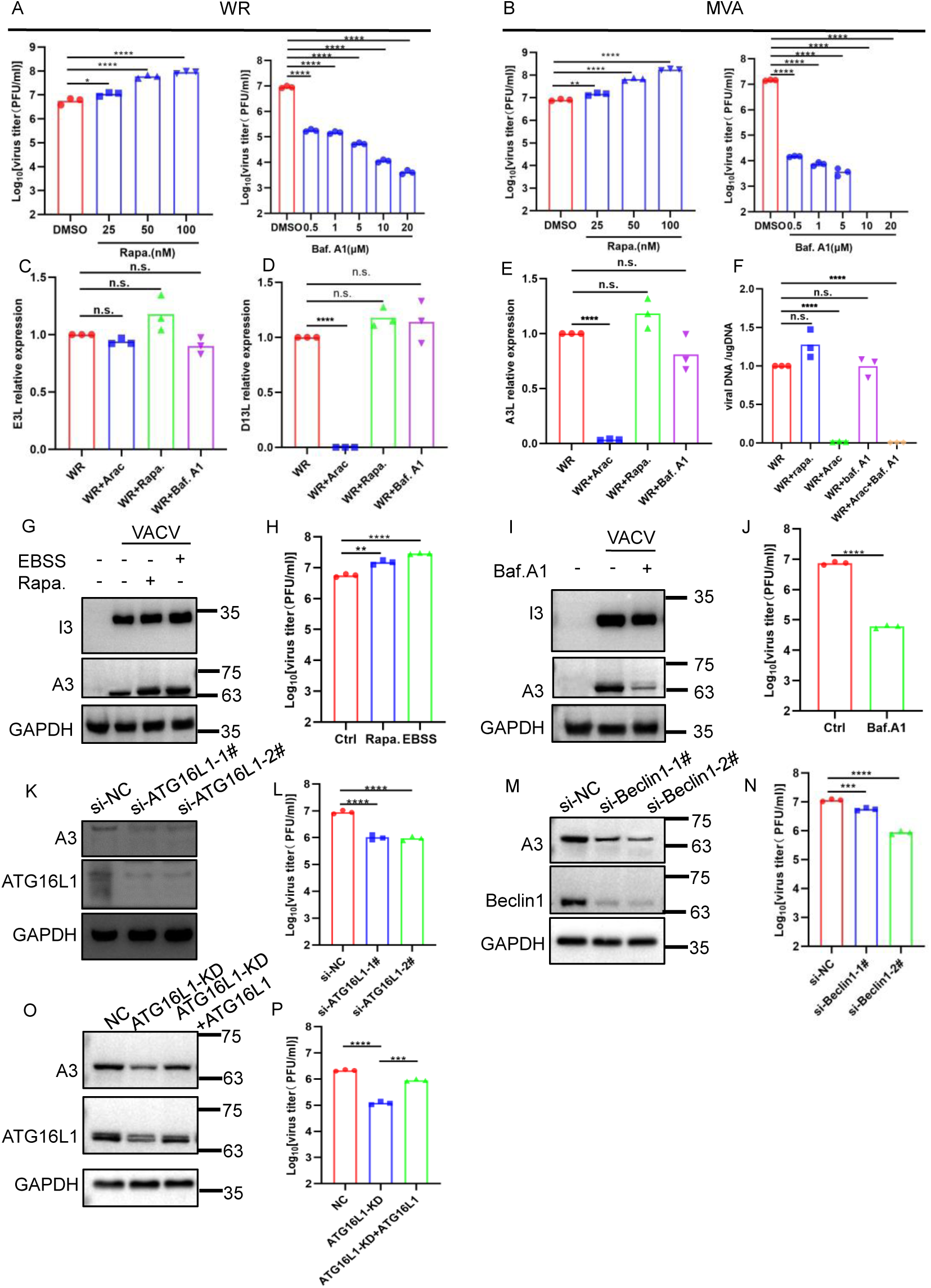
Prohibition of autophagy dampens VACV replication. Human A549 (A) or DF-1 cells (B)were infected in triplicate with VACV-WR and MVA at an MOI of 3 in the presence of rapa. or baf. A1 at the indicated concentrations. Virus yield at 24 hpi was determined by a plaque assay on BS-C-1 or DF-1 cells. (C-F) A549 cells were infected with VACV-WR in the presence of AraC (40 mg/mL), rapa. (100nM) or baf. A1 (1μM) for 2 h or 8 h and total RNA and DNA were harvested for mRNA and viral genomic DNA quantification using quantitative realtime-PCR with VACV-specific primers. (G-J) A549 cells were left untreated, starved for 2 h with EBSS or infected with VACV-WR in the presence of rapa. (100nM) or baf. A1 (1μM). At 12 hpi, the synthesis of viral proteins in infected cells was examined by Western blotting analysis using the indicated antibodies (G and I), and virus titers were determined by a plaque assay on BS-C-1 cells (H and J). (K-N) A549 cells were transfected with si-NC as a negative control, siATG16L1 or siBeclin 1 for 48 h and were subsequently infected with VACV-WR at an MOI of 3. After 24 h, cell lysates were collected for Western blotting analysis with antibodies for A3, ATG16L1, Beclin 1 or GAPDH as a loading control (K and L), and virus titers were determined by a plaque assay described above (L and N). (O and P) WT A549 cells (NC), A549 cells stably expressing shATG16L1 (ATG16L1-KD) and A549 cells stably expressing shATG16L1 cells transfected with KD-resistant ATG16L1 (ATG16L1-KD+ATG16L1) were infected with VACV-WR at an MOI of 3. After 24 hpi, cells were lysed and proteins harvested and resolved by SDS-PAGE followed by Western blotting analysis with anti-ATG16L1 and anti-A3 antibodies (O). Virus titers were determined by a plaque assay (P). Data in A, B, C, D, E, F, H, J, L, N and P are shown as dots, and the bar represents the mean value. Data in A-P are representative of three independent experiments. Statistics: n.s., not significant, p>0.05; *P < 0.05; **P < 0.01; ***P < 0.001; ****P < 0.0001 by two-sided Student’s t test.

### 2.2. VACV infection leads to the activation of autophagy

We next aimed to investigate if VACV infection resulted in changes in cellular autophagic regulation. As the conversion of microtubule-associated protein 1A/1B-light chain 3-I (LC3-I) to LC3-II and the autophagic degradation of p62, a main cargo degraded by autolysosomes, are considered classic molecular markers for autophagic activation, we first monitored the status of autophagy by observing these two markers in A549 cells infected with VACV-WR or MVA at 3 PFU/cell. In both MVA and VACV-infected cells, the conversion from LC3-I to LC3-II presented an activation of autophagy upon infection since 8 hpi (Figures 2A and 2B). Interestingly, although MVA infection led to a discernable decrease of p62 (Figure 2A), the level of p62 in VACV-WR-infected cells was comparable to that of the mock cells, and remained unchanged throughout the infection, even in the presence of baf. A1 (Figure 2B). Moreover, the degradation of p62 in LSDV-infected cells was similar to that observed in VACV-WR-infected cells (Figure S2A). These data suggested VACV-WR and LSDV may possess a means to block the steps of autophagic flux required for p62 degradation, which is mysteriously absent for MVA.

**Figure 2.**
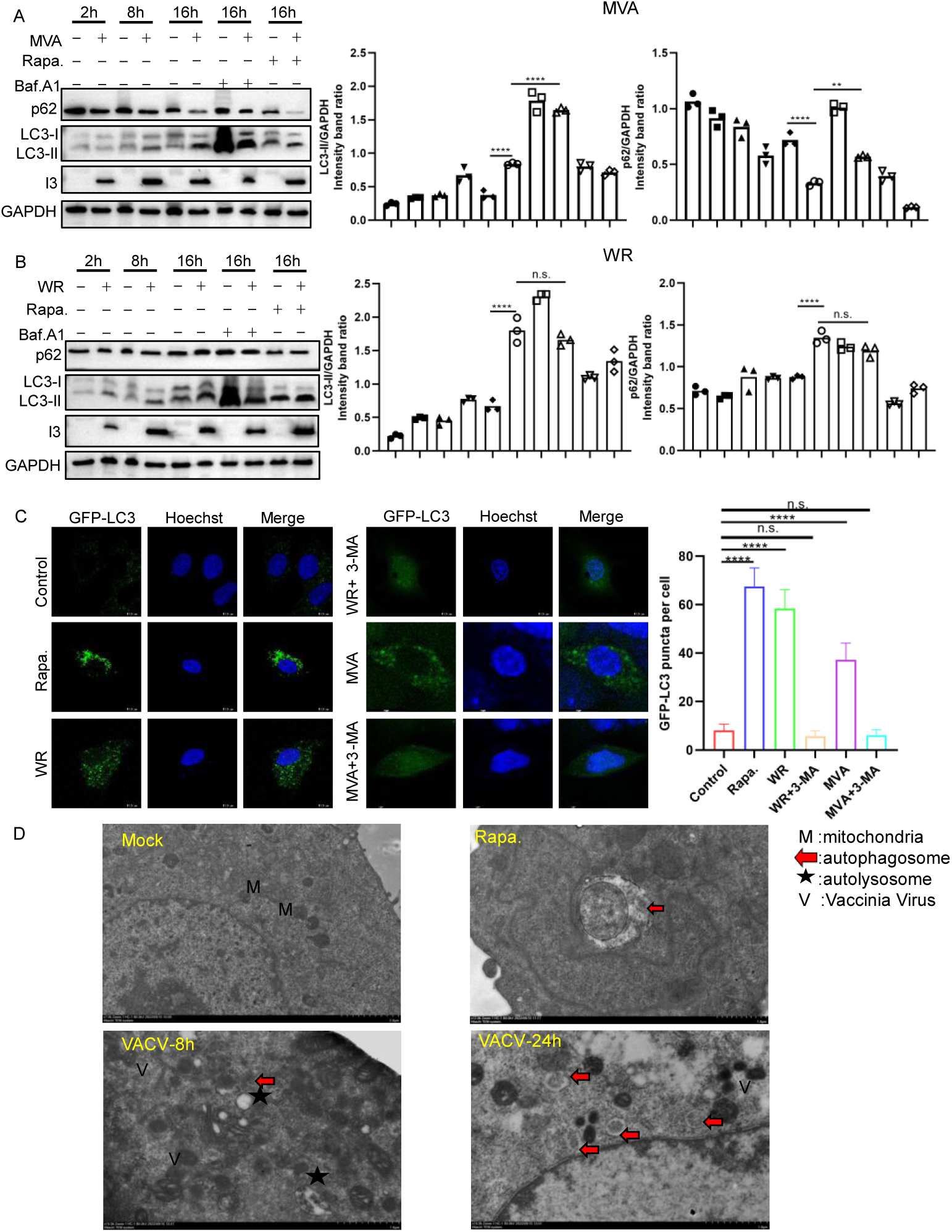
VACV infection promoted activation of autophagy. A549 were infected with VACV-WR (A) or MVA (B) at an MOI of 3 in the presence or absence of rapa. (100nM) or baf. A1 (1μM), and cell lysates were collected at indicated time points and subjected to Western blotting analysis with anti-p62, anti-LC3, anti-I3, and anti-GAPDH antibodies. (C) A549 cells stably expressing GFP-LC3 on coverslips were infected with VACV-WR, MVA at 3 PFU/cell or treated with 3-MA (1mM) or rapa. (100nM) and cells were fixed at 12 hpi, stained with Hoechst and observed with a fluorescent confocal microscope. Numbers of GFP-LC3 puncta (representing autophagosomes) were quantitated with Image J in 15 randomly-selected cells. Scale bars represent 10 μm. (D) A549 cells were mock-infected, treated with rapa. (100nM) or infected with VACV-WR at 3 MOI and were processed and analyzed for the accumulation of autophagosome via transmission electron microscopy. Red arrow indicates autophagic vacuoles, M indicates mitochondria, pentacle indicates autolysosome, and V indicates VACV. Data in A, B, and C represent the mean values of three independent experiments. Data in D are representative of three independent experiments. Statistics: n.s., not significant, p>0.05; ****P < 0.0001 by two-sided Student’s t test.

We next verified the observation by using an alternative assay, in which a GFP-labeled LC3 was stably expressed in A549 cells and the formation of green puncta in cells indicated autophagic activation. Although LC3 puncta was observed at a very low level in untreated control cells, the treatment of rapa., VACV-WR or VACV-MVA infection led to observable accumulation of LC3 puncta (Figure 2C), indicating activation of autophagy. A similar phenomenon was observed in A549 cells infected by LSDV (Figure S2B).

To directly visualize the formation of autophagosomes in infected cells, we employed transmission electron microscopy (TEM) to observe autophagosomes during VACV-WR infection. In mock-infected A549 cells, autophagic vacuoles were rarely observed (Figure 2D, top-left panel). However, in VACV-WR-infected A549 cells, fine structures of autophagosomes were observed at both 8 and 24 hpi, and more autophagosomes were observed at 24 hpi. (Figure 2D), suggesting that VACV infection resulted in the accumulation of autophagosomes. Taken together, these data demonstrated that VACV infection resulted in the activation of autophagy and the accumulation of autophagosomes. Importantly and surprisingly, while VACV-WR was able to inhibit the degradation of p62, MVA was unable to accomplish such inhibition.

### 2.3 VACV-WR, but not MVA, is able to block the fusion of autophagosome with lysosome

To further elucidate the difference between VACV-WR and MVA in autophagy regulation at a finer resolution, we employed an assay to differentiate complete and incomplete autophagy during virus infection. In this assay, a reporter plasmid containing a tandemly connected mCherry-GFP-LC3 was used to transfect A549 cells 24 h prior to VACV-WR or MVA infection. GFP is sensitive to and can be dampened in an environment with acidic pH, such as that in the lysosomes, whereas the mCherry is not^35^. When autophagosomes are not fused with lysosomes, which suggests incomplete autophagy, both green and red puncta can be observed and would appear as yellow puncta in merged images. In contrast, when autophagosomes fuse with lysosomes to complete autophagy, green puncta would rapidly disappear and only red puncta is observed (Figure 3A) ^35^. Cells transfected with the reporter plasmid mentioned above were infected with VACV-WR, MVA or LSDV at 3 PFU/cell and cells were fixed at 12 hpi and subjected to confocal microscopic analysis. In mock-infected cells, both green and red puncta were observed at a very low level (Figure 3B). In rapa. treated cells, the level of green signal remained low while red signal overwhelmed, suggesting the fusion of lysosomes with autophagosomes occured. However, when choloroquine (CQ) was added, the fusion of autophagosomes with lysosomes was inhibited as both green and red signals were detected and appeared as yellow puncta in merged images (Figure 3B), indicating incomplete autophagy. Importantly, in VACV-WR and LSDV-infected cells, both green and red signals were captured (Figures 3B and S3B), indicating incomplete autophagy. Interestingly and remarkably, in MVA-infected cells, the pattern of red/green signal was comparable to that of rapa.-treated cells, demonstrating the fusion of autophagosomes with lysosomes, which dampened the GFP signals (Figure 3B). We performed quantitation of the data by counting green and red puncta using Image J and the results were shown in Figure 3B. Moreover, we repeated this experiment in HeLa cells and observed very similar results (Figure S3A). Interestingly, in LSDV-infected HeLa and A549 cells, both situations were observed. In HeLa cells, which are non-permissive to LSDV, the loss of green signal was observed, showing similarity to MVA-infected cells. In contrast, in A549 cells, which are permissive to LSDV, we observed incomplete autophagy, manifested by the presence of both green and red signals (Figure S3B).

**Figure 3.**
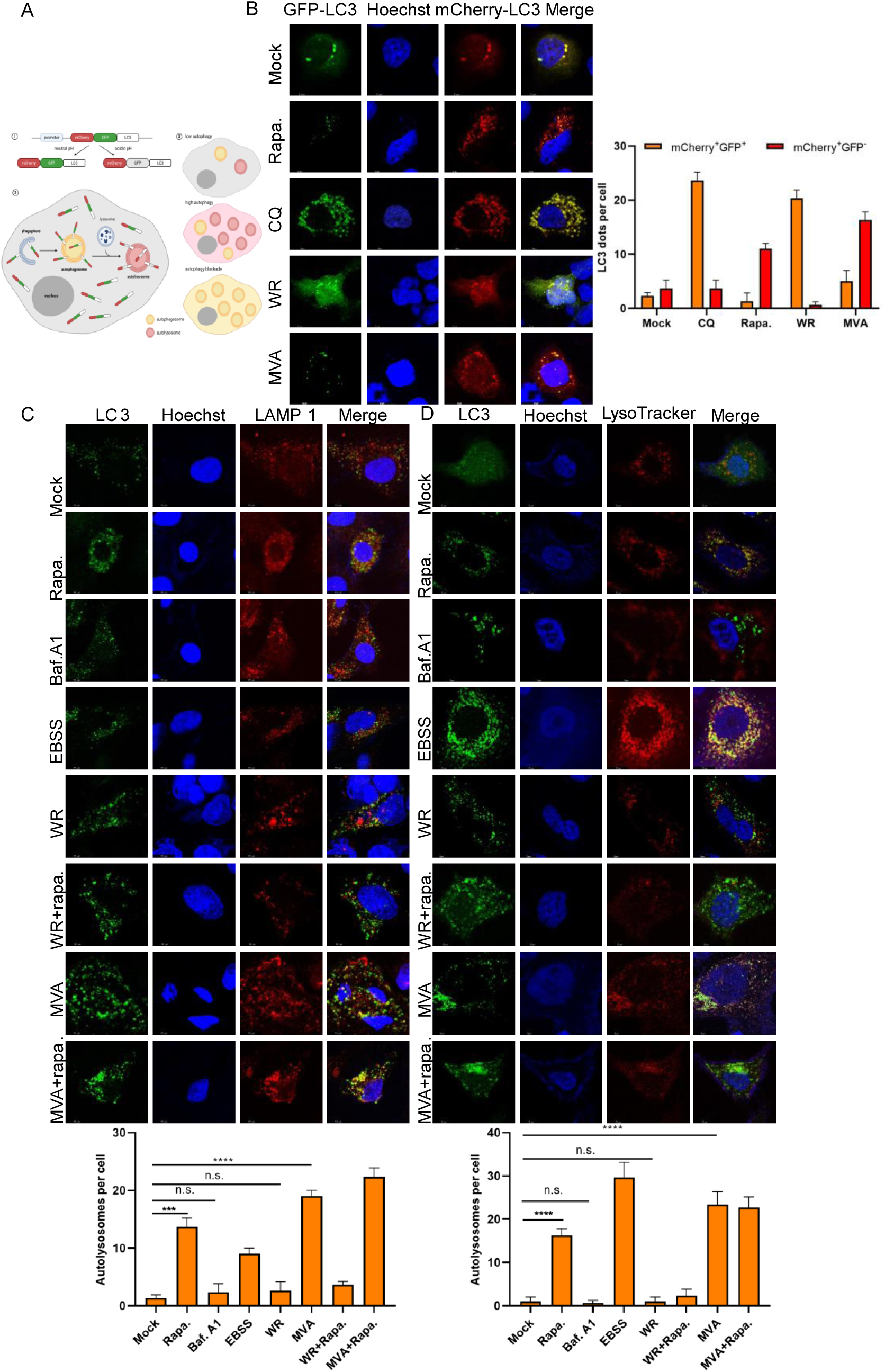
VACV-WR and MVA differed in their ability to inhibit autolysosome formation. (A) Diagram illustrating the working principle of the reporter plasmid mCherry-GFP-LC3. (B) A549 cells grown on coverslips were transfected with the mCherry-GFP-LC3 plasmid for 24 h and then were mock-infected, infected with VACV-WR or MVA at 3 PFU/cell, or treated with CQ (40μM) or rapa. (100nM) for 12 h. Cells were fixed, stained with Hoechst and analyzed with a fluorescent confocal microscope. Scale bars indicate 10 μm. Quantification of green and red puncta was performed with Image J in 15 randomly-selected cells. (C) A549 cells stably expressing LC3 on cover slips were infected with VACV-WR, MVA at 3 PFU/cell or treated with baf. A1 (5μM) or rapa. (100nM). At 12 hpi, cells were then fixed, permeabilized, blocked, and stained with primary antibodies to LAMP1 and followed by fluorescent conjugated secondary antibodies. Hoechst was used to stain nucleus. Scale bars represent 10 μm. (D) A549 cells stably expressing LC3 were infected with VACV-WR, MVA at 3 PFU/cell or treated with baf. A1 or rapa. as described above. Lysotracker was added to live cells to stain lysosomes. Images were taken as decribed above and quantification of colocalization was analyzed by Image J software in 15 randomly selected cells. Data in B, C, and D represent the mean values ± SD of three independent experiments. Statistics: n.s., not significant, p>0.05; *P < 0.05; ***P < 0.001; ****P < 0.0001 by two-sided Student’s t test.

We next investigated the co-localization of lysosomes and autophagosomes, which is a prerequisite for the fusion of autophagosomes with lysosomes. A549 cells that stably express LC3 were treated with the indicated drugs and subsequently infected with VACV-WR or MVA for 12 h prior to fixation and staining. LAMP1 was used to stain lysosomes. In agreement with our previous observation, rapa. and EBSS, which are autophagy agonists, promoted colocalization of autophagosomes and lysosomes (Figure 3C). The same phenomenon was observed in MVA-infected cells (Figure 3C). However, in VACV-WR and LSDV-infected cells, lysosomes were not found to be colocalized with LC3 (Figure 3C and S3C), indicating the blockage of the fusion of the two. These observations were verified in live cells in which lysotracker (Figure 3D), a cell permeable lysosome marker, was used to label the presence of lysosomes.

Taken together, our results demonstrated that VACV-WR and LSDV showed the capability to block the fusion of autophagosomes with lysosomes while MVA was unable to exert such a blockage. In addition, LSDV’s ability to block the fusion of autophagosomes and lysosomes was correlated with its capability to replicate in different cell lines. These findings led us to hypothesize that VACV-WR may encode a gene(s), whose product(s) may be responsible for blocking autophagosome-lysosome fusion, which was lost in MVA during its repeated passages in CEF cells. The loss of such genes in MVA may contribute to MVA’s host range restriction.

### 2.4 VACV A52 inhibits autophagosome-lysosome fusion

To identify the gene(s) responsible for the inhibition of the fusion, we compared the genomes of VACV-WR and MVA for the identification of candidate genes. As point mutations were found throughout the genome in MVA in comparison to VACV-WR, we decided to prioritize the genes that are either completely absent or those that exhibit more than 50% deletions. A total of 20 genes were found to be either completely absent or more than 50% deleted in MVA in comparison to VACV-WR. These genes were codon-optimized, synthesized and then cloned into a mammalian cell expression vector pcDNA3.1. As not all genes cloned were expressed successfully, only those that expressed effectively were further investigated. To screen for the inhibitor for autophagosome-lysosome fusion, A549 cells that stably express LC3-GFP were transfected with the above genes and then infected with MVA at 3 PFU/cell. The infected cells were then fixed and observed directly for GFP or stained with an anti-LAMP1 antibody. A positive target would be a viral protein that disrupts the colocalization of LC3-GFP and LAMP1, as observed in VACV-WR-infected cells (Figure 4A and S4A). In addition, all candidate genes were screened for their ability to block p62 degradation upon MVA infection (Figures S4B and S4C). Using these assays, we screened 20 VACV proteins and successfully identified WR178 (A52) as a positive candidate. The results showed A52 was able to disturb the colocalization of LC3 and LAMP1 observed in MVA-infected cells (Figure 4A). Moreover, although MVA infection led to degradation of p62, transfection of A52 was able to avert this phenomenon in a dose-dependent manner (Figure 4B).

**Figure 4.**
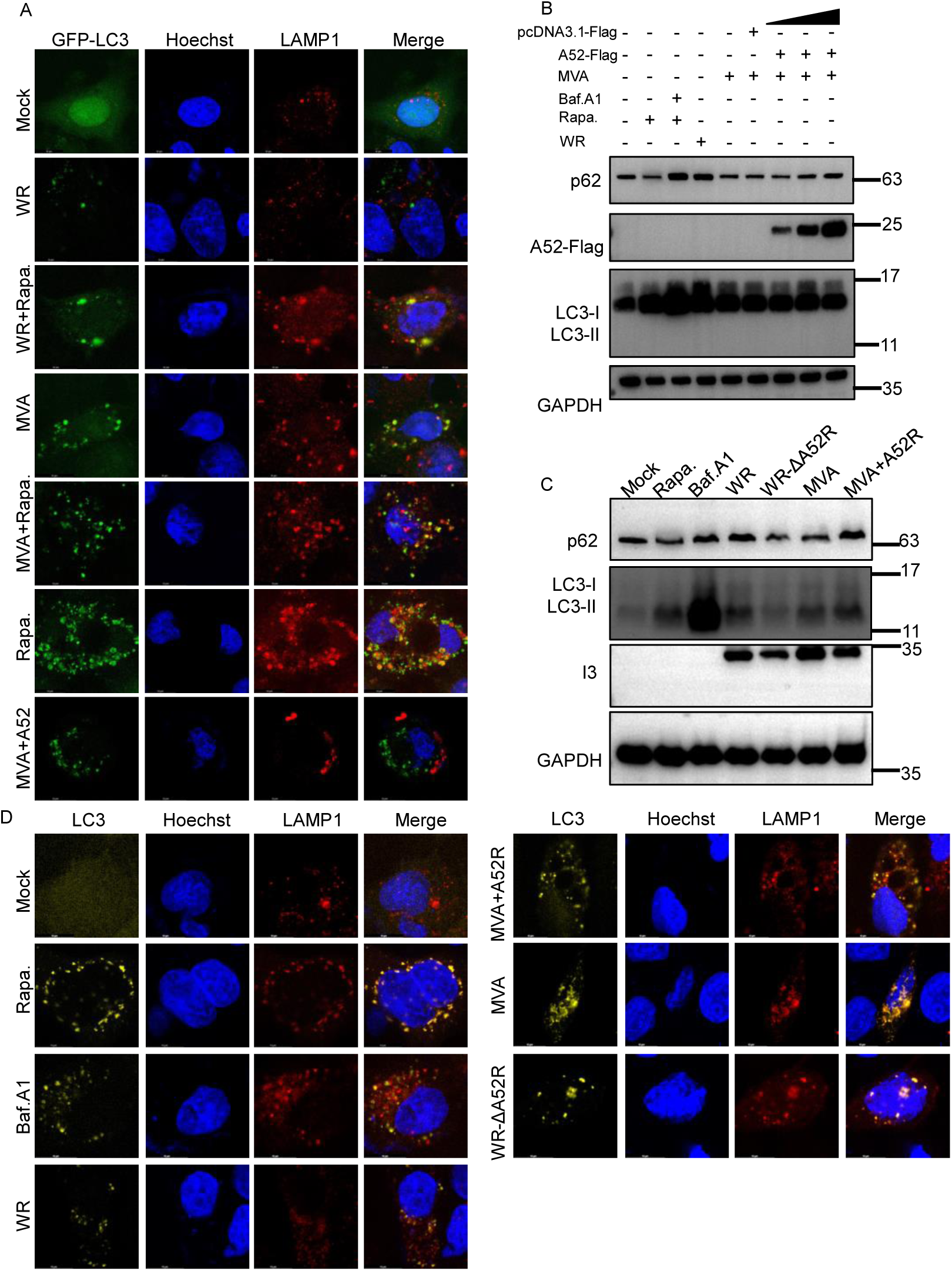
The effect of A52 on the fusion of autophagosome with lysosome. (A) A549 cells stably expressing LC3-GFP were left untreated, infected with VACV-WR or MVA at 3 MOI in the presence or absence of rapa. (100nM), or transiently transfected with a plasmid encoding a Flagged tagged A52 (1500 ng) prior to MVA infection. At 12 hpi, cells were then fixed, permeabilized, blocked, and stained with primary antibodies to LAMP1 and followed by fluorescent conjugated secondary antibodies. Hoechst was used to stain nucleus. Scale bars represent 10 μm. (B) Human A549 cells were transfected with a vector enconding A52-Flag at concentrations of 0.1, 0.5, and 1.5 μg/mL for 24 h and then infected with MVA at 3 PFU/cell for 12 h. VACV-WR-infected cells and cells treated with baf. A1 and rapa. were included as controls. Cell lysates were analyzed by SDS-PAGE followed by Western blotting analysis with anti-p62, anti-Flag, anti-LC3B, or anti-GAPDH antibodies. (C) Human A549 cells were infected with VACV-WR, VACV-WR-ΔA52R, MVA or MVA+A52R at 3 PFU/cell or treated with baf. A1 (1μM) or rapa. (100nM) for 12 h. Cell lysates were analyzed by SDS-PAGE and Western blotting with anti-p62, anti-I3, anti-LC3B, or anti-GAPDH antibodies. (D) A549 cells were infected with VACV-WR, VACV-WR-ΔA52R, MVA, MVA+A52R at 3 PFU/cell or treated with baf. A1 (5μM) or rapa. (100nM) for 12 h. Cells were then fixed, permeabilized, blocked, and stained with primary antibodies to LAMP1 or LC3 and followed by fluorescent conjugated secondary antibodies. Hoechst was used to stain nucleus. Scale bars represent 10 μm. Data in A-D are representative of three independent experiments.

To further investigate the biological function of A52, we replaced the open reading frame (ORF) of A52R in VACV-WR with a cassette containing a GFP to generate a recombinant virus with deleted A52R, and the virus was named VACV-WR-ΔA52R. In addition, a p11-driven-A52 ORF was inserted into the intergenic region between ORF069 and ORF070 in MVA to generate a recombinant MVA expressing A52, which was designated MVA+A52R. Both recombinant viruses were plaque-purified and successful deletion or insertion was confirmed with Sanger sequencing. Experiments described above were performed to monitor the autophagic degradation of p62 and co-localization between LC3 and lysosomes upon infection with these recombinant viruses. Compared to VACV-WR, the level of p62 was much lower in cells infected with VACV-WR-ΔA52R, indicating VACV-WR-ΔA52R’s ability to block p62 degradation was compromised by depleting A52R. Moreover, insertion of A52R into MVA was able to increase the level of p62 upon MVA’s infection (Figure 4C), suggesting A52 enabled MVA to inhibit p62 degradation.

Next, we observed the subcellular localization of endogenous LC3 and lysosomes in A549 cells infected with VACV-WR, VACV-WR-ΔA52R, MVA or MVA+A52R. The results indicated that A52 was essential for disrupting the colocalization between endogenous LC3 and lysosomes as both VACV-WR-ΔA52R and MVA were unable to complete such disruption (Figure 4D). In summary, our results demonstrated that VACV A52 was responsible for the inhibition of the fusion of autophagosomes with lysosomes.

### 2.5 A52 interacts with human SNAP29

Next, we aimed to elucidate the molecular mechanism by which A52 functions to block the fusion of autophagosomes with lysosomes. Synaptosomal-associated protein 29 (SNAP29) is a member of the SNARE (soluble NSF attachment protein receptor) family and is a pivotal factor that mediates the fusion of autophagosomes with lysosomes. Through interacting with other SNARE proteins such as syntaxin17 (STX17) and VAMP8, SNAP29 brings the membranes of the autophagosome and lysosome into close proximity for the fusion to occur. We hypothesized that A52 blocked the fusion through interacting with SNAP29. To test this hypothesis, the interaction between A52 and SNAP29 was first examined by a co-immunoprecipitation (co-IP) assay using ectopically expressed A52 and SNAP29. A52 was detected in the protein complex pulled down with antibodies for SNAP29-Myc (Figure 5A) and the reciprocal co-IP (Figure 5B) also revealed a positive association between the two proteins. To minimize artifacts, A52 protein expressed by the recombinant virus MVA+A52R was employed as a bait for endogenous SNAP29 in A549 cells. Co-IP analysis confirmed that the endogenous SNAP29 was indeed associated with viral A52 (Figure 5C). Next, we attempted to verify if A52 colocalized with SNAP29 during virus infection. A549 cells were infected with MVA+A52R at 3 PFU/cell and cells were fixed at 12 hpi and stained for A52 with anti-Flag antibody and endogenous SNAP29 with a monoclonal antibody for human SNAP29. Both proteins were observed in the cytoplasm, and the merge image showed a colocalization pattern of the two (Figure 5D).

**Figure 5.**
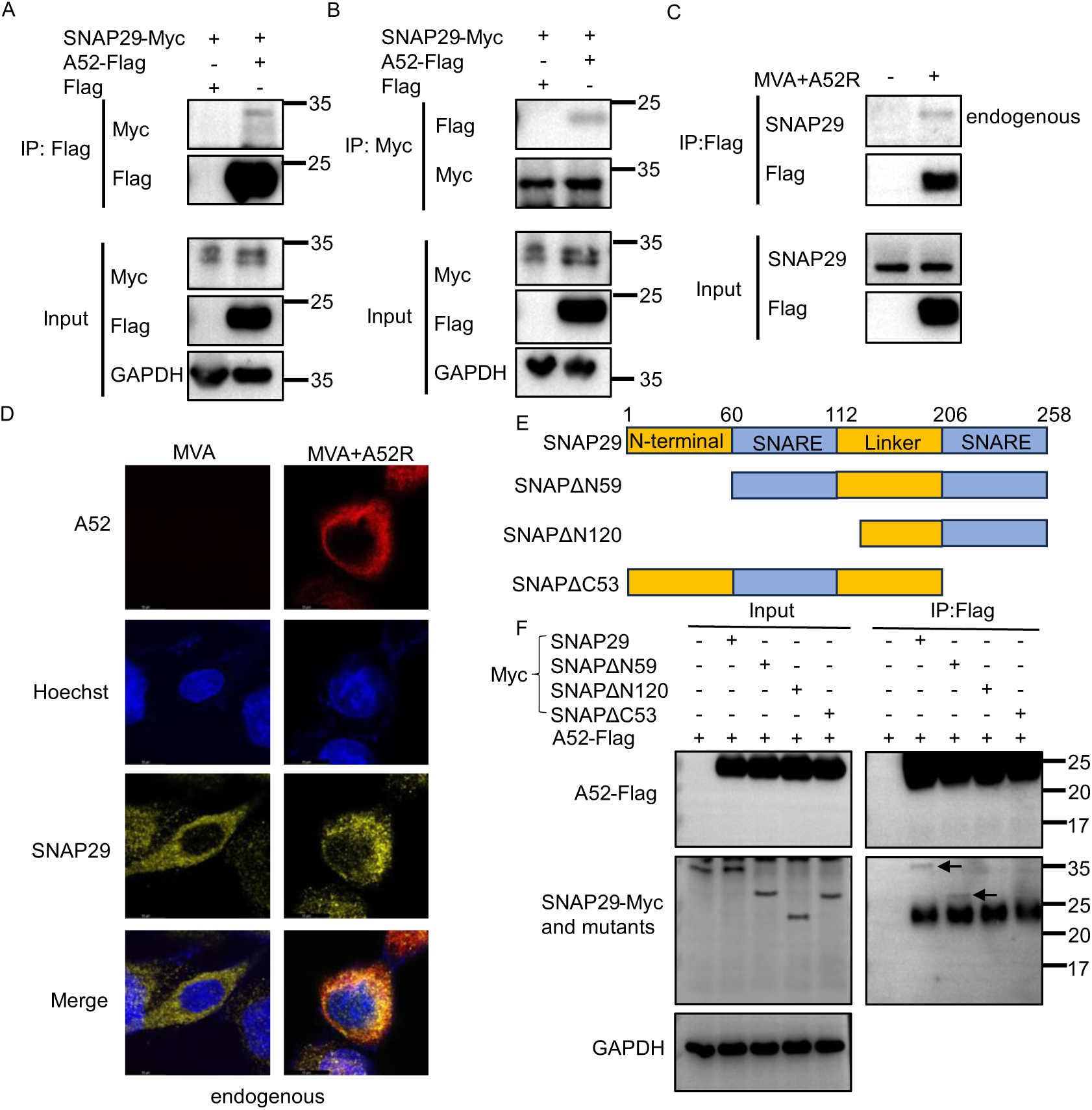
A52 is associated with human SNAP29. Human A549 cells were co-transfected with plasmids encoding Myc-tagged SNAP29 and Flag-tagged A52 for 36 h. Cell lysates were pre-cleared with control magnetic beads and then incubated with Flag-conjugated (A) or Myc-conjugated (B) beads at 4°C for 18 h. Beads were extensively washed and proteins were eluted with SDS loading buffer and analyzed by SDS-PAGE and Western blotting described above. (C) A549 cells were mock-infected or infected with MVA+A52R at 3 MOI for 24 h. Cell lysates were pre-cleared with control magnetic beads or Flag-conjugated beads at 4°C for 18 h. Beads were extensively washed and proteins were eluted with SDS loading buffer and analyzed by SDS-PAGE and Western blotting with anti-SNAP29, anti-Flag antibodies. (D) A549 cells grown on coverslips were infected with MVA or MVA+A52R at 3 MOI for 12h, and then fixied, stained with antibodies for SNAP29, Myc-tag and Hoechst for 2 h and then incubated with fluorescent secondary antibodies for 1 h. Images were taken with a fluorescent confocal miscroscope. Scale bars represent10 μm. (E) A series of SNAP29 truncation mutants were constructed and illustrated. (F) A549 cells were co-transfected with vectors encoding Myc-tagged SNAP29 or its mutants, and a Flag-tagged A52 for 36 h. Cell lysates were pre-cleared with control magnetic beads or Flag-conjugated beads at 4°C for 18 h followed by extensive washing. Proteins were eluted with SDS loading buffer and resolved by SDS-PAGE followed by Western blotting analysis using primary antibodies for Flag, SNAP29 and GAPDH. Data in A, B, C, D, and F are representative of three independent experiments.

Human SNAP29 contains two SNARE motifs (aa 60-112 and aa 206-258) divided by a linker region ^36^. To map the region responsible for its interaction with A52, we constructed three SNAP29 truncation mutants (Figure 5E) and examined their interaction with A52 using a co-IP analysis described above. The results showed that deletion of either of the SNARE motifs from SNAP29 completely sabotaged its interaction with A52, while removal of the N-terminal region had no such effect (Figure 5F). Altogether, these data demonstrated SNAP29 associated with viral A52 through its SNARE motifs.

### 2.6 A52 blocks Autophagosome-Lysosome Fusion by jeopardizing the interaction between SNAP29 and its binding partners STX17 and VAMP8

The formation of autolysosomes, a critical step in autophagy, is a delicately regulated process that involves the interplay among several cellular components including SNAP29, STX17, VAMP8 and VPS39. SNAP29 serves as an adaptor that binds to STX17 on autophagosomes and VAMP8 on lysosomes, which brings the two vesicles into close proximity for fusion to occur. To investigate if A52 blocks autolysosome formation by compromising SNAP29’s interaction with STX17 and VAMP8, the association between the two pairs was examined in the presence and absence of viral A52 by co-IP analysis. Although ectopic expression of SNAP29 was capable of pulling down STX17 in A549 cells, this interaction was mitigated by the presence of A52 (Figure 6A). Reciprocal co-IP using STX17 as the bait exhibited similar results (Figure 6B). In addition, results from the confocal microscopic analysis using transfected STX17 and SNAP29 exhibited that the existence of A52 abolished the colocalization between the former two proteins (Figure 6C). To verify if A52 expressed from VACV-WR was able to perform the same function, A549 cells co-transfected with SNAP29-Myc and STX17 were infected with VACV-WR or VACV-WR-ΔA52R and the association between SNAP29 and STX17 was determined by a co-IP analysis described above. When baiting with SNAP29, the amount of STX17 co-IPed was greatly reduced when cells were infected with VACV-WR. However, the reduction was much less prominent in VACV-WR-ΔA52R-infected cells (Figure 6D). The difference observed was not due to any difference in virus input as the abundance of viral I3 was comparable between the cells infected with VACV-WR or VACV-WR-ΔA52R. As SNAP29 also binds to VAMP8, we next examined the role of A52 in interfering this interaction. As shown in Figure 6E, the presence of A52 evidently attenuated the association between SNAP29 and VAMP8, suggesting A52 was able to hamper the interaction of SNAP29 with both STX17 and VAMP8. The orthologs of A52 from MPXV and LSDV exhibited only 24% and 22.07% identity to that of VACV (Figure S5). To explore if they were able to obstruct SNAP29’s biological function in human cells, orthologs of A52R from MPXV and LSDV were synthesized and cloned into a mammalian expression vector. A co-IP analysis described above was performed. Surprisingly, although displaying low amino acid sequence identities to VACV A52, both orthologs were able to hinder the interaction between human SNAP29 and STX17 or VAMP8 tested in the co-IP analysis (Figures S6A-S6D).

**Figure 6.**
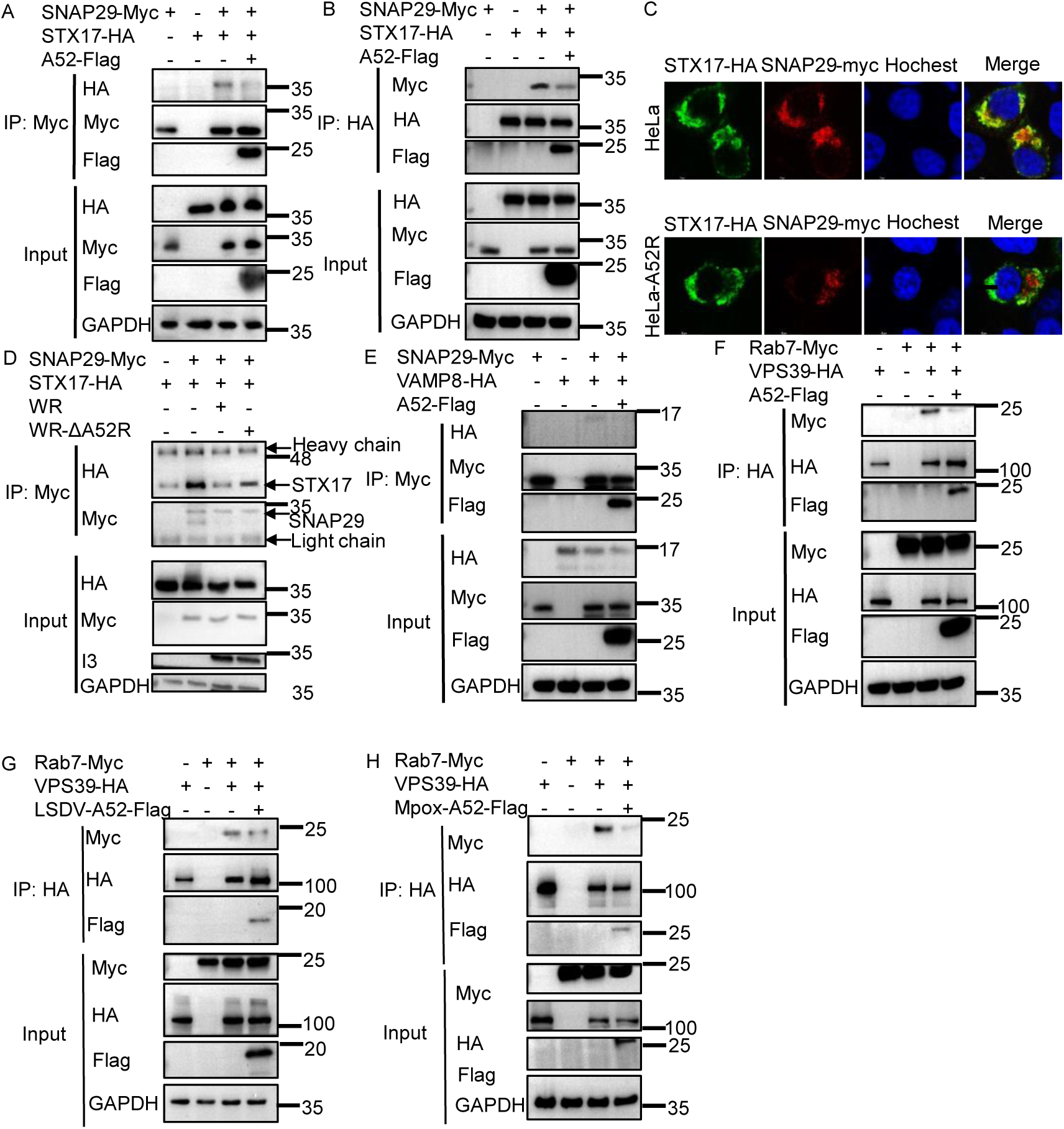
A52 hampered the interaction between SNAP29 and its binding partners STX17 and VAMP8. Human A549 cells were co-transfected with plasmids encoding Myc-tagged SNAP29, HA-tagged STX17, and Flag-tagged A52 for 36 h. Cell lysates were pre-cleared with control magnetic beads and then with Myc- (A) or HA-conjugated (B) beads at 4°C for 18 h. Beads were extensively washed and proteins were eluted with SDS loading buffer and resolved by SDS-PAGE and Western blotting analysis. (C) A549 cells on coverslips were co-transfected with SNAP29-Myc, STX17-HA and A52-Flag for 24 h. Cells were then fixed, stained with anti-Myc, anti-HA antibodies and Hoechst and images were taken with a fluorescent confocal microscope. Scale bars represent 10 μm. (D) A549 cells were co-transfected with SNAP29-Myc and STX17-HA for 24 h, and then infected with VACV-WR or VACV-WR-ΔA52R for 12 h. Cell lysates were precleared with control magnetic beads and then incubated with Myc-conjugated beads at 4°C for 18 h. Beads were extensively washed and proteins were eluted with SDS loading buffer and resolved by SDS-PAGE and Western blotting analysis. (E) A549 cells were co-transfected with SNAP29-Myc, VAMP8-HA, and A52-Flag for 36 h. Cell lysates were processed as described in (D). (F-H) A549 cells were co-transfected with Rab7-Myc, VPS39-HA and Flag tagged A52 from VACV, LSDV or MPXV. Cell lysates were pre-cleared with control magnetic beads and then incubated with HA-conjugated beads at 4°C for 18 h. Beads were extensively washed and proteins were eluted with SDS loading buffer and resolved by SDS-PAGE and Western blotting analysis. Data in A-H are representative of three independent experiments.

Apart from the SNARE complex, homotypic fusion and protein sorting (HOPS) complex is also known to mediate the fusion of autophagosomes with lysosomes. To analyze if A52 jeopardizes the HOPS complex during autophagy, the interaction between VPS39 and rab7 was resolved in the presence and absence of A52 using co-IP analysis. As shown in Figure 6F, the interaction between the two HOPS components was weakened when A52 was ectopically expressed. Interestingly, A52 orthologs from LSDV and MPXV exhibited a similar potency in such a process (Figures 6G and 6H). These data collectively demonstrated that A52 from VACV, MPXV and LSDV jeopardized the formation of the assembly of the SNARE and HOPS complexes required for the fusion of autophagosomes with lysosomes.

### 2.7 VACV-A52 promotes the degradation of SNAP29 via the proteasome pathway

As A52 was capable of compromising the interaction between SNAP29 and its binding partners STX17 and VAMP8 during autophagy. We were curious to inquire if A52 impacts the stability of SNAP29 during viral infection. For this purpose, the abundance of cellular endogenous SNAP29 was determined by Western blotting analysis when increasing amounts of VACV-A52 or LSDV-A52 were transfected into A549 cells. The endogenous levels of SNAP29 decreased as the amount of VACV-A52 increased (Figure 7A), but remained relatively steady when LSDV-A52 was transfected (Figure 7B). A time-dependent analysis also showed the reduction of SNAP29 in response to only VACV A52, but not LSDV A52 (Figures 7C and 7D). To inspect if SNAP29 was reduced when A52 is expressed by virus, A549 cells were infected with VACV-WR, VACV-WR-ΔA52R, MVA or MVA+A52R and the level of endogenous SNAP29 was determined by Western blotting analysis. As shown in Figure 7E, the amount of SNAP29 observed correlated with the presence of A52 and was more abundant in the cells infected with viruses lacking A52R (Figure 7E).

**Figure 7.**
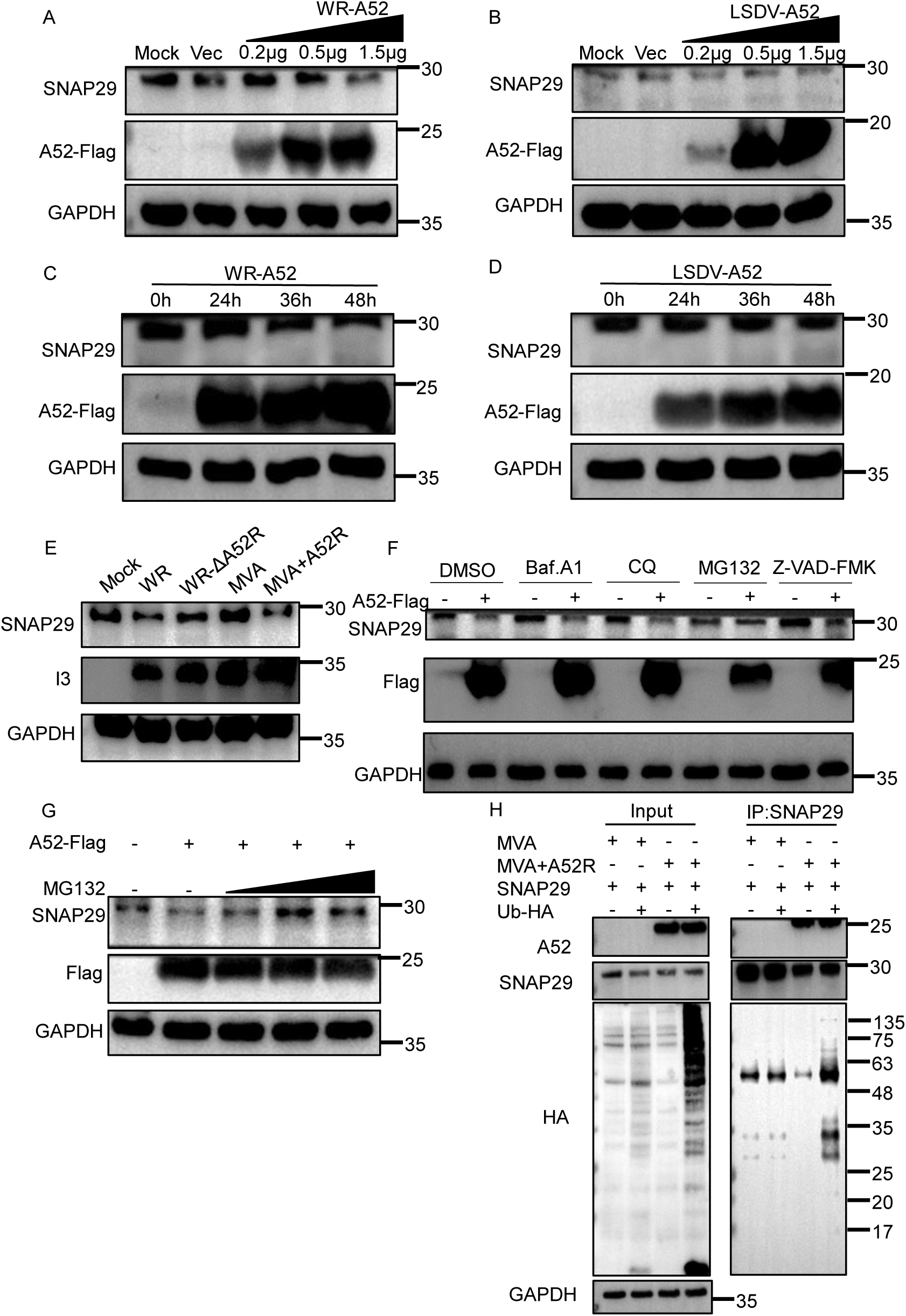
VACV-A52 facilitated proteasome-mediated SNAP29 degradation. A549 cells were transfected with vectors expressing VACV-WR-A52 (A) or LSDV-A52 (B) at concentrations of 0.2, 0.5, and 1.5 μg/mL for 36 h. Cell lysates were resolved by SDS-PAGE followed by Western blotting analysis with anti-SNAP29, anti-Flag, anti-GAPDH primary antibodies. A549 cells were transfected with plasmids encoding A52 from VACV (C) or LSDV (D) at 1.5 μg/mL and total proteins were collected at 0, 24, 36, and 48 h post transfection. Protein samples were resolved with SDS-PAGE and Western blotting analysis with primary antibodies for human SNAP29, Flag or GAPDH. (E) A549 cells were infected with VACV-WR, VACV-WR-ΔA52R, MVA or MVA+A52R for 24 h, and cell lysates were resolved by SDS-PAGE followed by Western blotting analysis with anti-SNAP29, anti-I3, anti-GAPDH antibodies. (F) A549 cells were transfected with the A52-Flag or empty vector plasmids for 24 h, and then the cells were treated with Baf. A1 (1 μM), CQ (40 μM), MG132 (5 μM) or Z-VAD-FMK (10 μM) for 12 h. DMSO was included as a negative control. Western blotting analysis was conducted to detect endogenous SNAP29 synthesis using protocols described above. (G) A549 cells were transfected with A52-Flag for 24 h in the presence or absence of MG132 at various concentrations. After 12 h, cell lysates were analyzed by SDS-PAGE and Western blotting analysis using anti-SNAP29, anti-Flag, anti-GAPDH antibodies. (H) A549 cells were infected with MVA or MVA+A52R at 3 PFU/cell. After 2 h, cells were washed with PBS and then transfected with an empty vector or vector enconding HA-tagged Ubiquitin for 36 h. Cell lysates were then precleared with control magnetic beads and then incubated with SNAP29-conjugated beads at 4°C for 18 h. Beads were extensively washed and proteins were eluted with SDS-loading buffer and resolved by SDS-PAGE followed by Western blotting analysis with primary antibodies for Flag (A52), SNAP29 or HA (ubiquitin). Data in A-H are representative of three independent experiments.

To investigate the nature of the reduction of SNAP29, A549 cells were transfected with a Myc-tagged A52 in the presence or absence of a series of chemical inhibitors targeting different protein degradation pathways, including baf. A1, CQ, MG132 and Z-VAD-FMK. Total protein lysates were collected 36 h post transfection and the level of SNAP29 was resolved by Western blotting analysis using a monoclonal antibody for endogenous SNAP29. In agreement with the previous observation, introduction of A52 resulted in an observable reduction of endogenous SNAP29. While the treatment of baf. A1, CQ and Z-VAD-FMK had minimal effect on the level of SNAP29, the addition of MG132 remarkably ameliorated the depression of SNAP29 caused by A52 (Figure 7F), insinuating that the reduction of SNAP29 might be the result of protein degradation via the proteasomal pathway. A dose-dependent experiment was performed and the results further confirmed that the addition of MG132 led to the rescue of A52-mediated SNAP29 degradation (Figure 7G). To further corroborate this conclusion, the ubiquitination of SNAP29 was inspected in cells infected with MVA or MVA+A52R (Figure 7H). Compared with MVA, the ubiquitination level of SNAP29 was greater in cells infected with MVA+A52R (Figure 7H), suggesting A52 induced the ubiquitination of SNAP29. These results demonstrated that A52 enhanced the ubiquitination of SNAP29 and the subsequent degradation via a proteasome-dependent mechanism.

### 2.8 SNAP29 is a host restriction factor for VACV lacking A52

MVA is replication-deficient in the majority of mammalian cells and almost all human cells^6^. Previous reports illustrated that zinc-finger antiviral protein (ZAP) and FAM111A contributed to the restriction of MVA in human cells. As our data demonstrated that VACV-WR was capable of blocking autophagosome-lysosome fusion while MVA was unable to do so, we hypothesized that the inability to block the formation of autolysosome might contribute to the restriction of MVA in human cells. To test this hypothesis, we manipulated the synthesis of SNAP29 by small interfering RNA (siRNA) and examined the replication of MVA and recombinant MVA+A52R in these cells compared to control cells. Transfection of si-SNAP29 led to distinct reduction of SNAP29 protein in A549 cells, and resulted in an increase of MVA titers up to 10× fold. Nevertheless, the effect of SNAP29 depression was much less prominent for the replication of MVA+A52R as the recombinant expressing A52 already showed a 10× fold enhancement in virus replication (Figures 8A, 8B and 8C). To ensure the effect on virus replication was due to SNAP29 but not the off-target effect of siSNAP29, A549 cells were transfected with SNAP29 prior to infection with MVA or MVA+A52R and viral titers were monitored. Ectopic expression of SNAP29 greatly reduced the level of MVA (∼15× fold) but had a much less prominent effect on MVA+A52R (∼3× fold) (Figures 8D, 8E and 8F).

**Figure 8.**
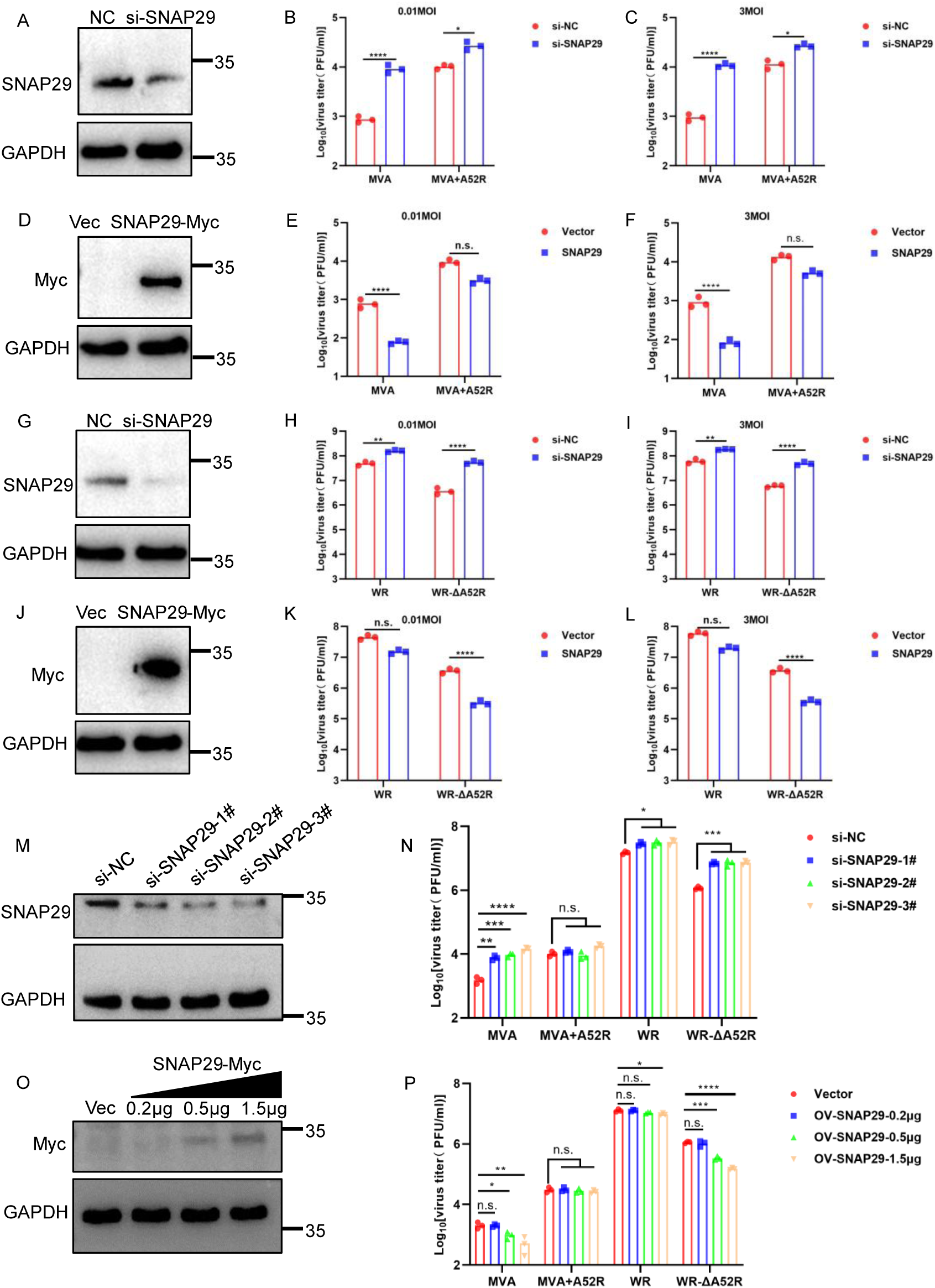
SNAP29 inhibits the replication of MVA in A549 cells. Human A549 cells were transfected with siNC or siSNAP29 (A) for 48 h and then infected in triplicates with MVA or MVA+A52R at 0.01 (B) or 3 (C) PFU/cell. Viruses were harvested at 24 (MOI=3) or 48 hpi (MOI=0.01) and virus titers were determined by plaque assay on DF-1 cells. Human A549 cells were transfected with SNAP29-Myc (D) for 24 h and then infected in triplicates with MVA or MVA+A52R at 0.01 (E) or 3 PFU/cell (F). Viruses were collected at 48 or 24 hpi and titers were determined by plaque assay on DF-1 cells. Human A549 cells were transfected with siNC or siSNAP29 (G) for 48 h and then infected in triplicates with VACV-WR or VACV-WR-ΔA52R at 0.01 (H) or 3 PFU/cell (I). Viruses were collected at 24 or 48 hpi and virus titers were determined by plaque assay on BS-C-1 cells. Human A549 cells were transfected with SNAP29-Myc (J) for 24 h and then infected in triplicates with MVA or MVA+A52R at 0.01 (K) or 3 PFU/cell (L). Viruses were collected at 48 or 24 hpi and virus titers were determined by plaque assay on BS-C-1 cells. (M) A549 cells were transfected with three different pairs of siSNAP29 for 48 h and cell lysates were resolved by SDS-PAGE followed by Western blotting analysis with primary antibodies to SNAP29 and GAPDH. (N) Cells transfected with siNC or siSNAP29 described in (M) were infected in triplicates with MVA, MVA+A52R, VACV-WR or VACV-WR-ΔA52R at 3 PFU/cell for 24 h and virus titers were determined by plaque assay on DF1 (MVA) or BS-C-1 (WR) cells. (O) A549 cells were transfected with SNAP29-Myc at concentrations of 0.2, 0.5, or 1.5 μg/mL for 24 h and cell lysates were resolved by SDS-PAGE followed by Western blotting analysis with primary antibodies to SNAP29 and GAPDH. (P) Cells transfected with increasing concentrations of SNAP29 described in (O) were infected in triplicates with MVA, MVA+A52R, VACV-WR or VACV-WR-ΔA52R at 3 PFU/cell for 24 h and virus titers were determined by plaque assay on DF1 or BS-C-1. Data in B, C, E, F, H, I, K, L, N and P are shown as dots, and the bar represents the mean value. Data in A-P are representative of three independent experiments. Statistics: n.s., not significant, p>0.05; *P < 0.05; **P < 0.01; ***P < 0.001; ****P < 0.0001 by two-sided Student’s t test.

Next, we analyzed the consequence of SNAP29 suppression on the replication of VACV-WR and VACV-WR-ΔA52R. The loss of A52R resulted in a 10× fold decline in virus titer, but the reduction was rescued by suppressing SNAP29 expression via siSNAP29 (Figures 8G, 8H and 8I). In addition, ectopically expressed SNAP29 caused a 15× fold decrease of the titers of VACV-WR-ΔA52R but had an insignificant impact on the replication of VACV-WR (Figures 8J, 8K and 8L).

To further consolidate our conclusion, we used three pairs of small interfering RNA (siRNA) targeting SNAP29 to knock down SNAP29 and examined the replication of MVA and recombinant MVA+A52R or WR and VACV-WR-ΔA52R in these cells compared to control cells. Transfection of si-SNAP29 led to distinct reduction of SNAP29 protein in A549 cells (Figure 8M), and resulted in an increase of MVA and WR-ΔA52R titers up to 10× and 8× folds (Figure 8N). Nevertheless, the effect of SNAP29 depression was much less prominent for the replication of MVA+A52R and VACV-WR (Figure 8N). In addition, Transfection of SNAP29-Myc led to a reduction of titers of MVA and WR-ΔA52R in a dose-dependent manner, but had an insignificant impact on the replication of MVA+A52 and VACV-WR (Figures 8O and 8P). Taken together, these data suggested that SNAP29 served as a restriction factor for MVA in human cells, which could be overcome by VACV A52.

## 3. DISCUSSION

Autophagy is a fundamental cellular process responsible for maintaining cellular homeostasis by degrading damaged proteins or invading pathogens^13^. Understanding how viruses manipulate autophagy can provide insights into the molecular mechanism of viral pathogenesis and host-virus interactions. Our data showed autophagy inhibition suppressed VACV’s replication and presented a molecular mechanism by which VACV A52 overcame the fusion of autophagosomes with lysosomes (summarized in Figure 9). A previous report indicated that the deletion of ATG5 and Beclin 1 from mouse embryonic fibroblasts (MEF) did not impact VACV’s replication^31^. A previous report indicated that Vaccinia virus subverts xenophagy through phosphorylation and nuclear targeting of p62 and nuclear translocation of p62 was dependent upon p62 NLS2 and correlated with VACV kinase B1 and F10 mediated phosphorylation of p62 T269/S272^37^. In this study, we demonstrated that pharmaceutical inhibition or the suppression of ATG16L1 significantly impaired the replication of VACV-WR, MVA and LSDV. Our observations did not necessarily contradict with the previous observations, as our analyses differed in two key aspects. First, our data were obtained using human cell lines, including A549 and HeLa, whereas the previous study utilized primary cells collected from mouse embryos. Variations in the autophagy machineries between human and mouse cells could influence virus-modulated autophagic dynamics. Secondly, the previous study employed a modified version of VACV-NYCBH (New York City Board of Health) strain ^31^, which served as the parental strain of VACV-WR. VACV-WR evolved from repeated passage of the NYCBH strain in various animals and a variety of cell lines. The heightened sensitivity of VACV-WR to autophagy manipulation may result from the virus’s adaptation to specific cellular environments.

**Figure 9.**
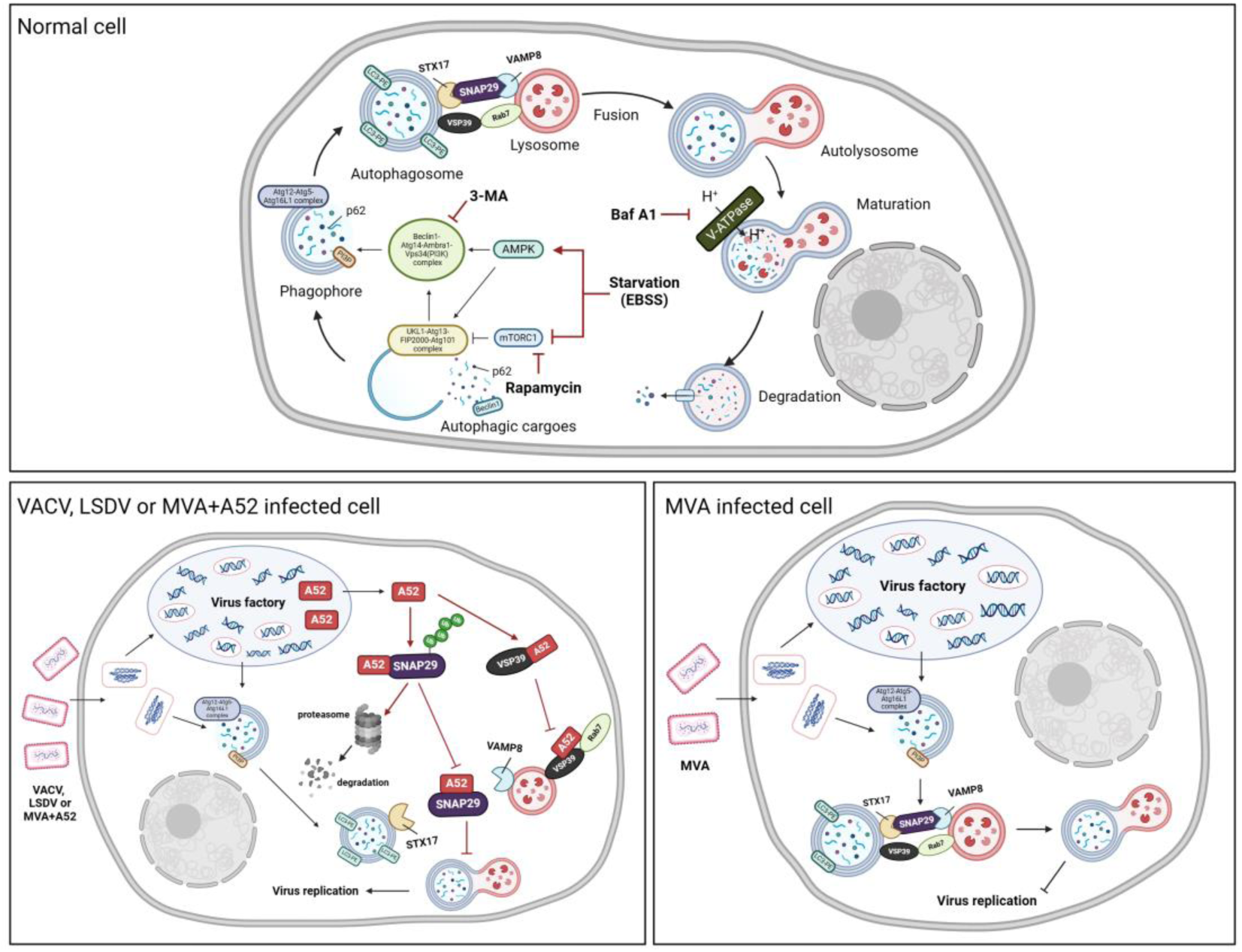
Mechanism of A52 in modulating the fusion of autophagosome and lysosome The figure was generated using Biorender.

Both chemical autophagy inhibitors and the depression of crucial autophagy factors severely hamper virus replication. We aimed to observe autophagosomes in VACV-WR-infected cells by TEM and a large number of autophagosomes were observed. Our data demonstrated that VACV-WR inhibited the fusion of autophagosomes and lysosomes by interfering with the SNARE complex (STX17-SNAP29-VAMP8) and HOPS complex (VPS39-Rab7), leading to the failure of cargo degradation and the accumulation of autophagosomes. The specific reason why VACV tends to halt autophagy at the point prior to the formation of autolysosomes is unknown. We hypothesized that this may prevent viral proteins from being degraded by autolysosomes, which requires further investigation to prove. Our analyses and others suggested that multiple viral proteins might be involved in regulating autophagy during viral infection. Identifying the viral proteins responsible for triggering autophagy would be an intriguing research topic in the future, and may provide target for attenuation of the VACV or other related poxviruses for vaccine development.

One of the most captivating observations was the phenotypic difference between VACV-WR and MVA in their ability to obstruct the fusion of autophagosomes with lysosomes. To delve into the molecular determinants behind, we cloned 20 genes absent in MVA but present in VACV-WR and conducted screening to assess their protein products’ capability to impede the formation of autolysosomes. An alternative approach was to delete genes from VACV-WR and select for mutants that lack the ability to inhibit the fusion. However, this approach carries risks, as there may be redundancy in the process, with multiple viral proteins independently capable of performing the inhibition. Indeed, our analyses revealed multiple viral proteins capable of inhibiting the fusion of autophagosomes with lysosomes, including A52, C10 and C11. Among which A52 exhibited the strongest capability and was selected for the subsequent research into the molecular mechanism. We also demonstrated that A52 was capable of working independently as MVA with an inserted A52R (MVA+A52R) showed potent inhibition of autolysosome formation comparable to that observed in VACV-WR-infected cells and the insertion of A52R greatly enhanced MVA’s replication in human cells. These results underscored the importance of blocking autolysosome-mediated degradation for efficient VACV replication.

The SNAP29 protein serves as a bridge that mediates the fusion of autophagosomes and lysosomes by connecting STX17 and VAMP8, anchored on autophagosomes and lysosomes, respectively ^36, 38^. VACV A52 was able to bind to SNAP29 through its SNARE motifs and such interaction effectively impaired the association between SNAP29 and its binding partners STX17 and VAMP8. These observations were consistent with the results reported by other groups using other viruses, such as Severe Acute Respiratory Syndrome Coronavirus 2 (SARS-CoV-2), Porcine reproductive and respiratory syndromevirus (PRRSV), and Human parainfluenza virus type 3 (HPIV3). As VACV A52 does not show any sequence similarity to other non-poxvirus proteins, these findings illustrate that various viruses have independently evolved a convergent mechanism to hinder autolysosome formation for their benefit. Importantly, A52 orthologs were found in 15 poxviruses from different genera, and we verified that the orthologs from mpox and LSDV were also capable of inhibiting the interaction between SNAP29 with STX17 and VAMP8, indicating that efficient blocking of autolysosome formation might be a conserved function in many poxviruses. In addition, we showed that A52 not only impaired SNAP29’s function, but also led to its proteasome-mediated degradation.

As poxviruses evolved strategies to inhibit SNAP29’s cellular function, we next asked if overexpressing SNAP29 was detrimental to virus replication. Our data exhibited that ectopic expression of SNAP29 showed inhibitory effect on virus replication for those viruses that lack A52R. Meanwhile, transient suppression of SNAP29 with RNAi enhanced the replication of VACV-ΔA52R or MVA but did not significantly impair the replication of VACV-WR or MVA+A52R. These results suggested that SNAP29 is a novel host restriction factor for MVA, and is antagonized by virus A52. In recent years, two host restriction factors for MVA have been identified, namely zinc-finger antiviral protein (ZAP) and FAM111A, which could be antagonized by VACV proteins C16 and C12, respectively ^9, 10^. Our discoveries added SNAP29 to this list and contributed fresh insights into comprehensive elucidation of the molecular mechanism underlying MVA’s inability to replicate efficiently in most mammalian cells.

Bioinformatic analyses revealed that A52 shows similarity to several other poxvirus BCL-2 proteins with immunomodulatory functions^39, 40^. Previous reports also illustrated that A52 was able to inhibit the activation of NF-κB and block the production of proinflammatory cytokines ^41–43^. Further research is needed to determine whether other BCL-2 proteins were able to antagonize the function of SNAP29. In addition, the fusion process of autophagosomes and lysosomes is tightly regulated by several factors in addition to SNAP29 ^44–52^. In fact, our study demonstrated that A52 also prevented the interaction between VPS39 and Rab7. Whether A52 can hijack other cellular proteins responsible for the fusion of autophagosomes with lysosomes needs further investigation. It is unclear whether the simultaneous interference of A52 with STX17-SNAP29-VAMP8 complex and VPS39-Rab7 complex is sufficient for prohibiting the fusion of autophagosomes with lysosomes. In addition, although we have observed that A52 plays a crucial role in blocking complete autophagy, other viral proteins or components may synergistically or independently promote this process to favor viral replication. Therefore, further research is needed to investigate the molecular interactions and functional effects between poxviruses and cellular autophagy.

## 4. Materials and methods

### Cell lines, plasmids and viruses

Alveolar type II-like epithelial (A549) cells, HeLa cells were purchased from Ningbo Ming Zhou bio CO., Ltd. MDBK [NBL-1] (cat#CL-0153), BHK-21[C-13] (cat#CL-0034), DF-1(cat#CL-0279) were kindly provided by Procell Life Science&Technology Co., Ltd. BS-C-1(cat#MZ-8048) cells were purchased from Ningbo Ming Zhou bio CO., Ltd. Establishment of A549 cells stably expressing GFP-LC3B was performed as follows. A549 cells were infected with Lentivirus containing GFP-LC3B (GV616, Genechem, Shanghai, China) for 4 h and then culture medium was removed and supplemented with fresh medium. Cells were then treated with 2.5μg/ml puromycin continuously for 7 days. The expression of LC3 was identified by WB analysis and fluoresecent confocal microscopic analysis. All cells were cultured at 37°C with 5% CO_2_ in Dulbecco’s modified Eagle’s medium (DMEM) supplemented with 10% (vol/vol) fetal bovine serum (FBS), 100 units of penicillin/ml, 100 µg/ml of streptomycin.

ORFs of SNAP29, STX17, VAMP8, VPS39 and Rab7 were cloned from A549 cDNA into a pcDNA3.1 vector. Genes from VACV-WR, MPXV and LSDV were codon-optimized and cloned into the pcDNA3.1 vector. Ub-HA plasmid was kindly provided by Junda Zhu and was previously described^10^.

VACV-WR and MVA were kindly provided by Bernard Moss from National Institutes of Health. LSDV-EGFP was constructed by Shijie Xie and reported previously^53^.

### Construction of recombinant MVA+A52R and VACV-WR-ΔA52R

Recombinant MVA+A52R was constructed by homologous recombination using fluorescent reporter genes (EGFP) for plaque selection. Briefly, a DNA cassette containing the A52R ORF regulated by an mH5 promoter followed by an EGFP ORF regulated by the P11 promoter was generated. The cassette containing mH5+A52R+P11+EGFP was inserted into the intergenic locus between MVA 069 and 070 by homologous recombination^8^. DF-1 cells were infected with parental MVA at 3 PFU/cell and then transfected with the DNA cassette described above. Viruses were harvested at 24 hpi and recombinant viruses were selected by selecting eGFP-positive clones. Positive clones were plaque-purified for 5× times in DF-1 cells. Viral DNA was isolated from the clones with the DNeasy Blood & Tissue kit (Qiagen) and sequences were verified by PCR followed by Sanger sequencing. VACV-WR-ΔA52R recombinant virus was generated as follows. A P11-directed eGFP sequence was cloned into a plasmid and flanked by 500 bp up- and downstream of the A52R ORF. The 1552-bp PCR product was inserted into pcDNA-3.1 vector and was used to transfect HeLa cells infected with VACV-WR at 3 PFU/cell for 2 h. A recombinant virus lacking A52 was isolated by selecting eGFP-positive virus clones and clonally purified by repeated plaque isolation. The loss of the A52 gene was confirmed by PCR and Sanger sequencing.

### Antibodies and chemicals

Rabbit anti-SNAP29 polyclonal antibody (12704-1-AP) was purchased from Proteintech. Rabbit anti-LC3 antibody (L7543) was purchased from Sigma. Rabbit anti-p62 (ab155686) was purchased from Abcam. Anti-LAMP1 antibody (#9091) was purchased from Cell Signaling Technology. Autophagy Antibody Sample Kit (4445T) was purchased from Cell Signaling Technology. Lyso-Tracker Red (C1046), anti-Flag mouse monoclonal antibody (AF2852), anti-Myc rabbit monoclonal antibody (AM933), anti-HA mouse monoclonal antibody (AF2858), BeyoMag™ Anti-Flag Magnetic Beads (P2115), BeyoMag™ Anti-HA Magnetic Beads (P2121) and BeyoMag™ Anti-Myc Magnetic Beads (P2118) were purchased from Beyotime. Rapamycin (Rapa.) (S1842) was purchased from Beyotime. Bafilomycin A1 (Baf. A1) (S1413) and 3-MA (3-Methyladenine) (S2767) were purchased from Selleck. Chloroquine diphosphate was kindly provided by Professor Kui Zhu from China Agricultural University.

### Virus infection and plaque assay

A549 or HeLa cells in 6-well plates were cultured to reach 90% confluency. Medium was removed and replaced with inoculum containing various viruses in complete DMEM supplemented with 2.5% FBS to reach a titer of 0.01 or 3 PFU/cell. After 2 h incubation at 37°C, the inoculum was removed, and cells were washed twice with 1× PBS and then supplemented with fresh medium. Total viruses were harvested at 48 hpi or 24 hpi by cell scaping and repeated free-thaw cycles. Virus titers were determined by plaque assay in BS-C-1 or DF-1 cells described previously^9^.

### Quantitative realtime-PCR

A549 cells in 6-well plates were infected with viruses indicated in figures at 3 PFU/cell for 1 h in triplicates. Cells were washed twice with 1× PBS and total DNA was extracted at 6 hpi with a DNeasy Blood/Tissue DNA mini kit (Qiagen) and total DNA yield was measured by a nanodrop spectrophotometer (Thermo Fisher Scientific). Equal amounts (1µg) of DNA from each sample were then serially diluted and subjected to quantitative realtime-PCR with gene-specific primers for VACV E11 and host GAPDH as the housekeeping control. The results of virus genomic DNA were normalized to that of GAPDH and relative DNA abundance was calculated and shown in figures. A549 cells in 6-well plates were infected with viruses at 3 PFU/cell in triplicates. Total RNA was extracted at 2 or 8 hpi with DNeasy Blood/Tissue DNA mini kit (Qiagen) and quantitated by a nanodrop spectrophotometer. Equal amounts (1µg) of RNA was reverse transcribed using a EasyScript^®^ First-Strand cDNA Synthesis SuperMix kit and subjected to quantitative realtime-PCR with gene-specific primers for virus E3, D13, A3 and host GAPDH as the housekeeping control.

### Transmission Electron Microscopy

Monolayer of HeLa cells was infected with VACV-WR or MVA at 3 PFU/cell for 8 or 24 h, or treated with rapa. (100nM) for 24 h. Trypsin digestion solutions were added to HeLa cells and then complete DMEM medium was added to terminate digestion. Cell suspension was collected into a 2 mL centrifugation tube and then centrifuged at 350 ×g for 5 min. Cell pellets were collected and washed with 1 mL 1× PBS for three times and then resuspended in 1 mL of 2.5% glutaraldehyde for fixation. Fixed samples were embedded in resin to create a solid block for sectioning. A ultramicrotome was employed to cut ultra-thin sections (around 50-100 nm) from the embedded sample. Transfer the ultra-thin sections onto a TEM grid and then apply heavy metal stains (e.g., uranyl acetate, lead citrate) to enhance contrast. The samples were then visualized using a transmission electron miscroscope (HITACHI HT7800).

### Confocal microscopy

Monolayers of A549 cells were grown on glass coverslips in 24-well plates and then infected with viruses indicated in figures 3B-3D, 4A, 4D, S4A, and S4D for the indicated time. Medium was removed and cells were fixed with ice-cold 4% paraformaldehyde for 20 min at room temperature, followed by incubation with 0.1% Triton X-100 for 20 min and blocked with 3% BSA diluted in 1× PBS for 30 min. LAMP1 antibody (1:200) and LC3 antibody (1:200) were added and incubated with cells at 4 °C overnight. Cells were then washed with 3% BSA for three times, followed by incubation with secondary antibodies conjugated with Alexa Fluor 488 or Alexa Fluor 555 or Alexa Fluor 647 at 37 °C in dark with agitation for 1 h. Hoechst was used to stain the nucleus for 5 min. Cells were washed multiple times before mounting on slides using ProLong Diamond Antifade reagent (Thermo Fisher Scientific). Images were taken on a Leica SP8 confocal microscope and images were processed with the Leica software (Leica Biosystems).

### Co-immunoprecipitation assays

Human A549 cells were co-transfected with vectors encoding Myc-tagged SNAP29, Rab7, HA-tagged STX17, VAMP8 and VPS39 and Flag-tagged A52 for 36 h. Cells were first washed once with ice-cold 1× PBS and lysed in wells with 150 µL IP lysis buffer with 1× PMSF (Beyotime Biotechnology) on ice. Protein lysates were collected and centrifuged for 10 min at 12,000 ×g at 4 °C and then precleared with control magnetic beads for 2h. Supernantant was collected and incubated with HA beads (Beyotime), Myc-Beads (Beyotime) or Flag-Beads (Beyotime) for overnight at 4 °C with rotation. The beads were then washed 6× times with 500μL 1× TBS buffer and the bound proteins were eluted by addition of 100μl 1× SDS-PAGE loading buffer containing 0.05M DTT and by boiling for 15 min. Supernatant was taken for SDS-PAGE followed by Western blotting analysis.

### Statistical analysis

All experiments were independently repeated for three times. The significance of the variability between different groups was determined by two-way ANOVA tests of variance using the GraphPad Prism software (version 6.0). P<0.05 was considered statistically significant, and P>0.05 was considered statistically nonsignificant.

## ACKNOWLEDGMENTS

We thank Dr. Bernard Moss from National Institutes of Health of USA for kindly providing MVA and VACV-WR. We thank the staff from the Life Science Experimental Center of CAU for facility use and help with microscopy and electron microscopy. We thank Prof. Rui Jia from Shandong University for his valuable advice.

## DECLARATION OF INTERESTS

No potential conflict of interest was reported by the author(s).

## FUNDING

This work was funded by the National Key Research and Development Program (2021YFD1800700), the National Natural Science Foundation of China (32172822), and the Beijing Nova Program (Z211100002121021) to C.P. The funders played no role in research design, data collection, analysis, the decision to publish or preparation of the manuscript.

---

**Figure S1.**
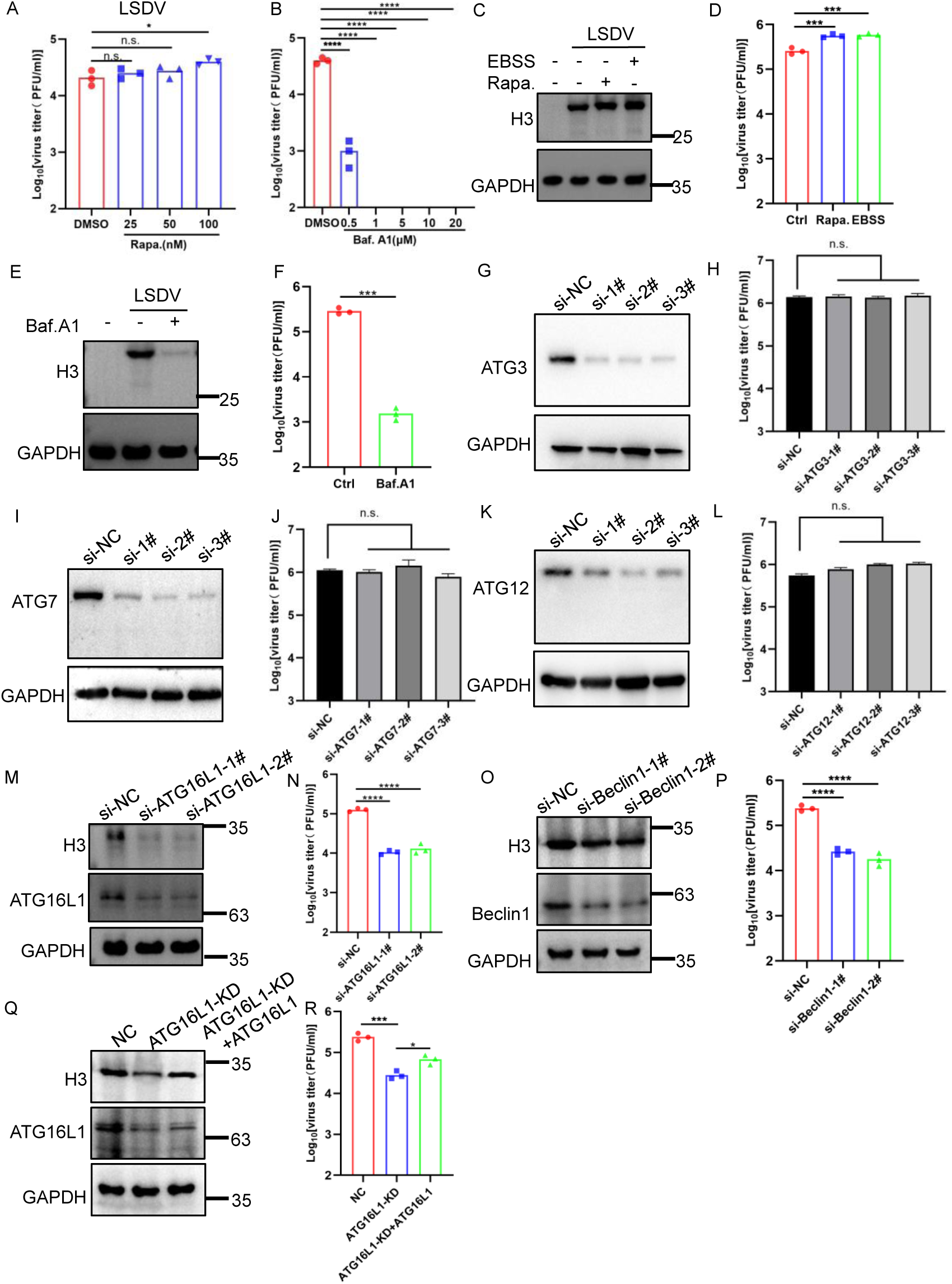
Autophagic inhibition suppresses LSDV’s replication and VACV-WR replication is independent of ATG3, ATG7, and ATG12. Human A549 were infected with LSDV at an MOI of 3 in the presence of rapa. (A) or baf. A1 (B) at the indicated concentrations. Virus yield at 24 hpi was determined by a plaque assay. (C-F) A549 cells were untreated, starved for 2 h with EBSS or infected with LSDV in the presence of rapa. (100nM) or baf. A1 (1μM). At 12 hpi, the expression of viral proteins in infected cells was detected by western blotting using the indicated antibodies (C and E), and virus titers were determined by a plaque assay (D and F). (G-L) A549 cells were transfected with si-NC, siATG3, si-ATG7, and si-ATG12 for 48 h and then were infected with the VACV-WR at an MOI of 3. After 24 hpi, cell lysates were collected for Western blot analysis (G, I, and K), and virus titers were determined by a plaque assay (H, J, and L). (M-P) A549 cells were transfected with si-NC, siATG16L1 or siBeclin 1, as indicated, for 48 h and then were infected with the LSDV at an MOI of 3. After 24 hpi, cell lysates were collected for Western blot analysis (M and O), and virus titers were determined by a plaque assay (N and P). (Q and R) A549 cells and A549 cells stably expressing shATG16L1 cells and A549 cells stably expressing shATG16L1 cells transfected with ATG16L1 were infected with LSDV at an MOI of 3. After 24 hpi, cells were lysed and proteins resolved by SDS-PAGE and Western blotting with anti-ATG16L1 and H3, GAPDH antibodies (Q), and virus titers were determined by a plaque assay(R). Data in A, B, D, F, H, J, L, N, P and R are shown as dots, and the bar represents the mean value. Data in A-R are representative of three independent experiments. Statistics: n.s., not significant, p>0.05; *P < 0.05; ***P < 0.001; ****P < 0.0001 by two-sided Student’s t test.

**Figure S2.**
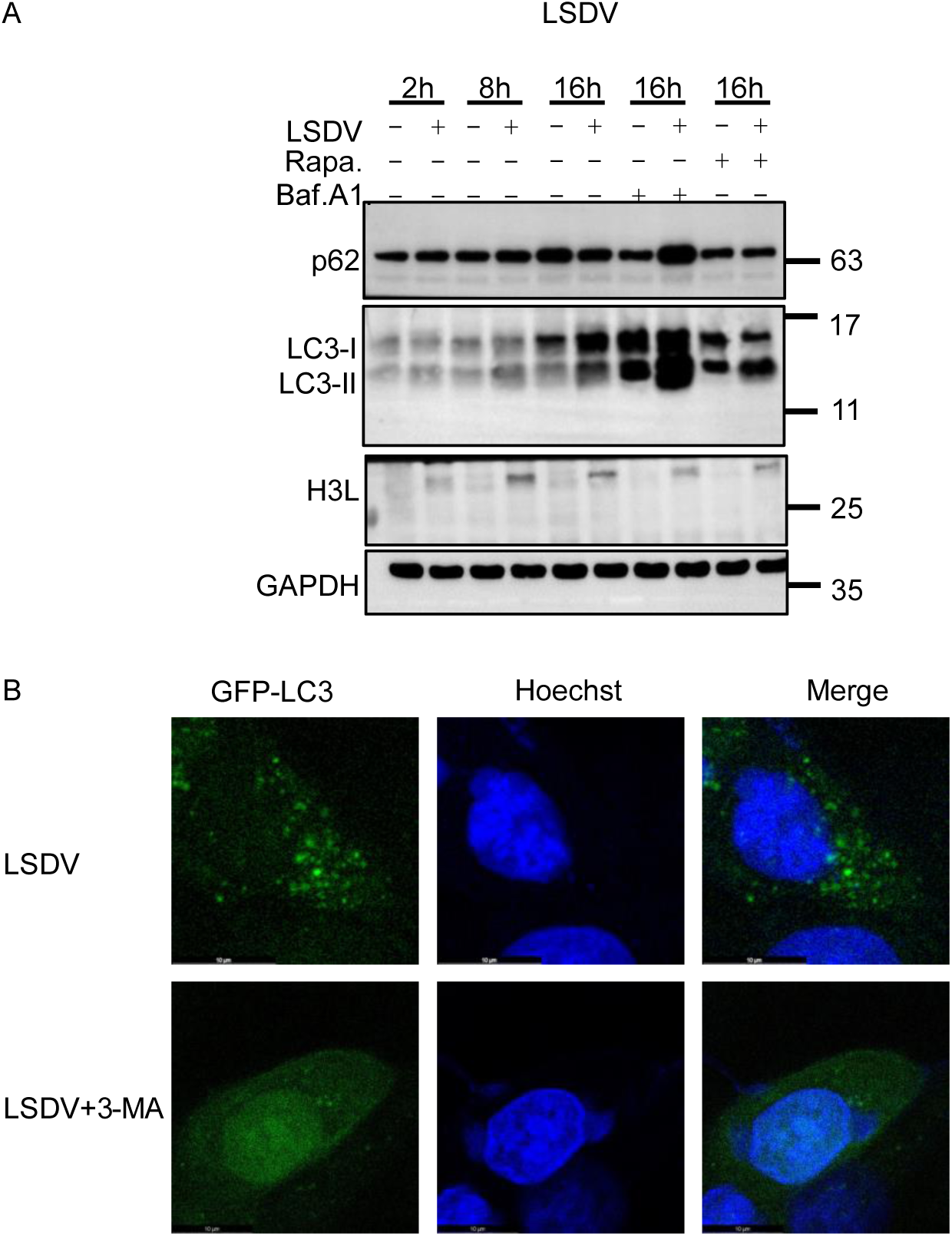
LSDV infection leads to the activation of autophagy. (A) A549 cells were infected with LSDV at an MOI of 3 in the presence or absence of rapa. (100nM) or baf. A1 (1μM), and cell lysates were collected at indicated time points and subjected to Western blot analysis with anti-p62, anti-LC3, anti-H3, and anti-GAPDH antibodies. (B) A549 cells stably expressing LC3 on coverslips were infected with LSDV at 3 PFU/cell or treated with 3-MA (1mM) or rapa. (100nM) and cells were fixed at 12 hpi, stained with Hoechst and observed with a fluorescent confocal microscope. Puncta of GFP-LC3 (representing autophagosomes) were observed. Scale bars, 10 μm. Data in A-B are representative of three independent experiments.

**Figure S3.**
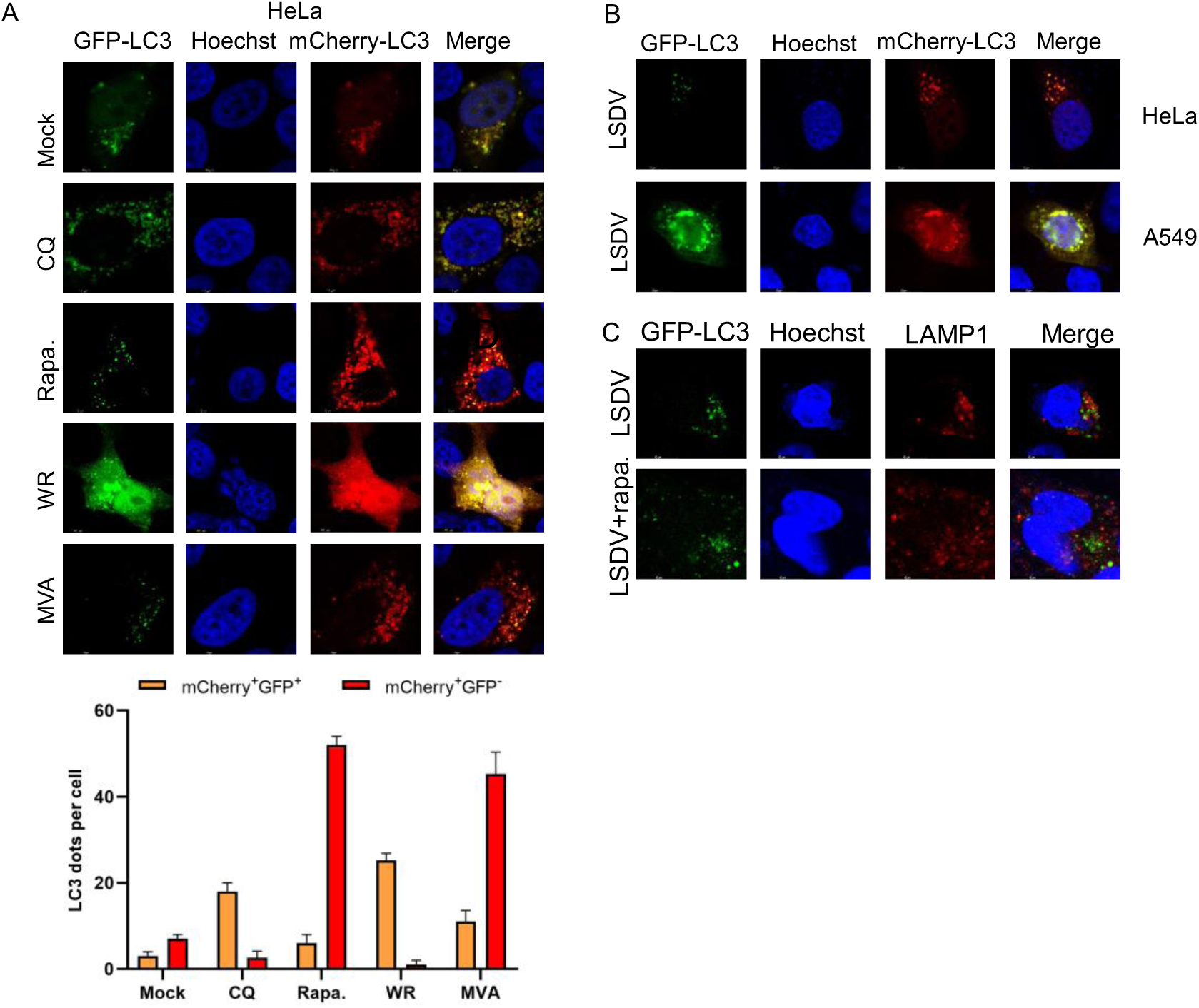
LSDV infection can block the fusion of autophagosome with lysosome in Permissive cell. (A) HeLa cells were transfected with mCherry-GFP-LC3 for 24 h and then were mock infected or infected with WR, MVA at 3 PFU/cell or treated with CQ or rapa. for 12 h, and then cells were fixed, permeabilized, blocked, stained with Hoechst and analyzed with a fluorescent confocal microscope. Scale bar, 10 μm. The graph shows the quantification of autophagosomes by taking the average number of dots in 15 cells. (B) HeLa or A549 cells were transfected with mCherry-GFP-LC3 for 24 h and then were mock infected or infected with LSDV at 3 PFU/cell and then cells were fixed, permeabilized, blocked, stained with Hoechst and analyzed with a fluorescent confocal microscope. Scale bar, 10 μm. (C) A549 cells stably expressing LC3 on cover slips were infected with LSDV at 3 PFU/cell or treated with rapa. (100nM). At 12 hpi, cells were then fixed, permeabilized, blocked, and stained with primary antibodies to LAMP1 and followed by fluorescent conjugated secondary antibodies. Hoechst was used to stain nucleus. Scale bars represent 10 μm. Data in A-C are representative of three independent experiments.

**Figure S4.**
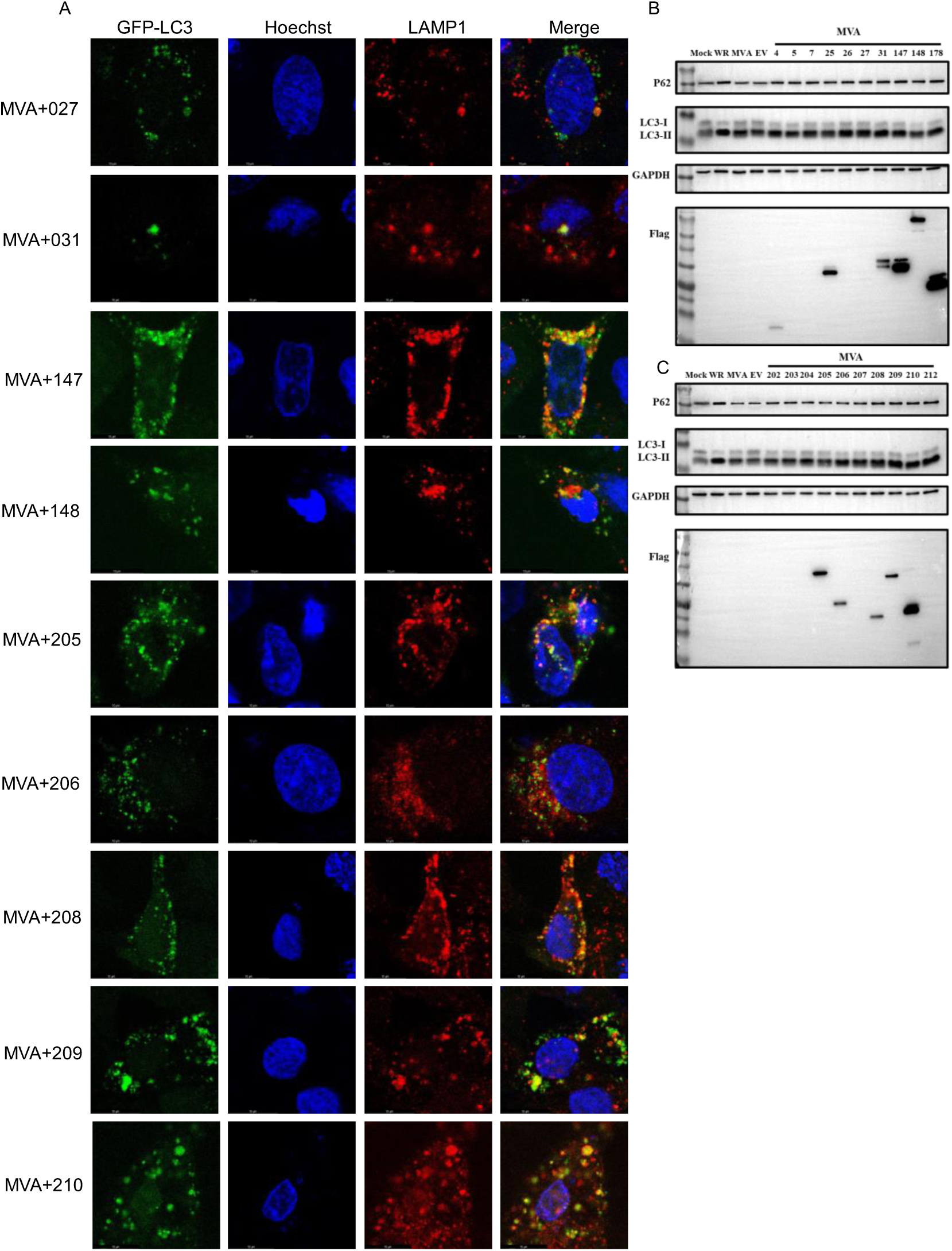
Identification of viral proteins responsible for inhibiting the fusion of autophagosomes and lysosomes. (A) Human A549 cells that stably express LC3-GFP were transfected with plasmids of all candidate genes prior to MVA infection at 3 PFU/cell. At 12 hpi, cells were fixed, permeabilized, blocked, and stained with primary antibodies to LAMP1 and followed by fluorescent conjugated secondary antibodies. Hoechst was used to stain nucleus. Scale bar, 10 μm. (B-C) Human A549 cells were infected with 3PFU/cell WR or MVA, after 2 h, cells were transfected with plasmids of all candidate genes, and cell lysates were collected for Western blot analysis with anti-p62, anti-LC3, anti-Flag, anti-GAPDH antibodies. Data in A-C are representative of three independent experiments.

**Figure S5.**
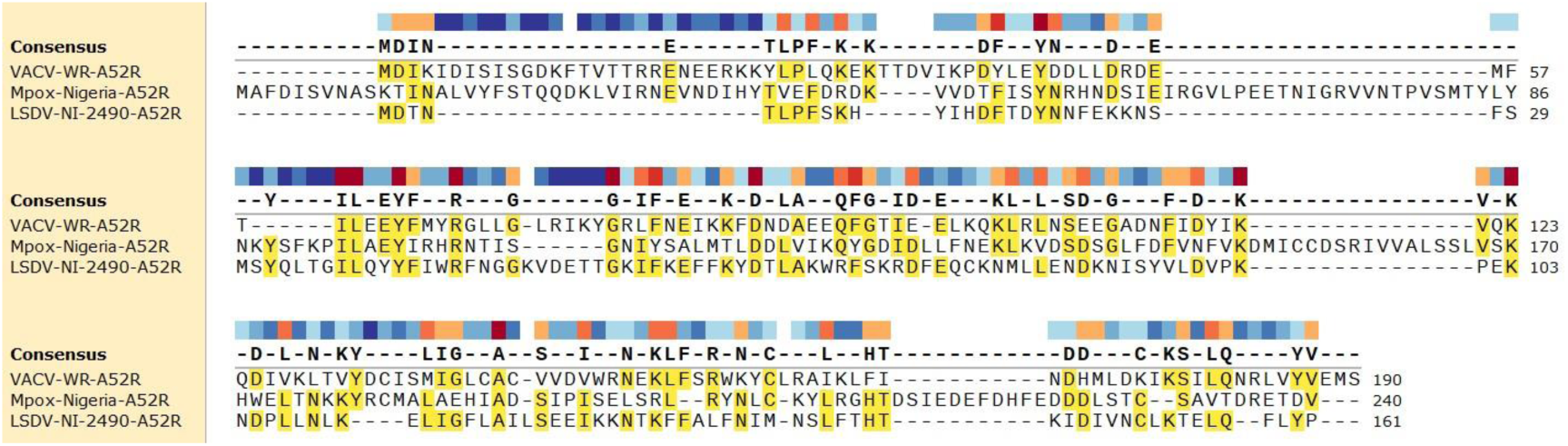
The orthologs of A52 from Mpox and LSDV exhibited 24% and 22.07% identity to that of VACV. The A52 amino acid sequences of VACV, Mpox, and LSDV were downloaded from NCBI (https://www.ncbi.nlm.nih.gov/), and snapgene was used for multiple sequence alignment.

**Figure S6.**
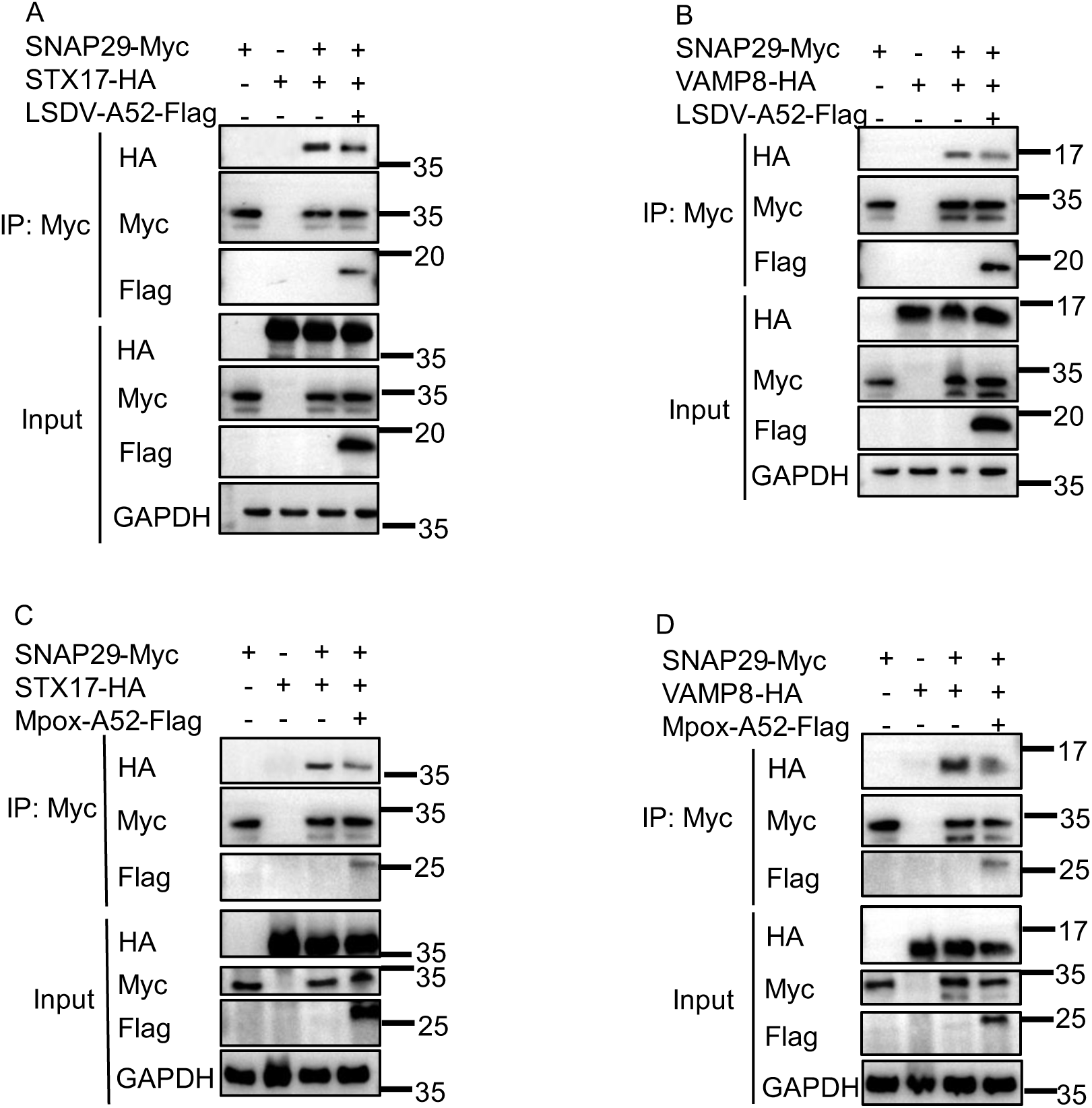
The orthologs of A52 from MPXV and LSDV block the interaction between SNAP29 and its binding partners STX17 and VAMP8. (A and C) Human A549 cells were co-transfected with SNAP29-Myc, STX17-HA, Flag tagged A52 from LSDV or MPXV for 36 h. Cell lysates were pre-cleared with control magnetic beads and then incubated with Myc-conjugated beads at 4°C for 18 h. Beads were extensively washed and proteins were eluted with SDS loading buffer and resolved by SDS-PAGE and Western blotting analysis. (B and D) Human A549 cells were co-transfected with SNAP29-Myc, VAMP8-HA, Flag tagged A52 from LSDV or MPXV for 36 h. Cell lysates were pre-cleared with control magnetic beads and then incubated with Myc-conjugated beads at 4°C for 18 h. Beads were extensively washed and proteins were eluted with SDS loading buffer and resolved by SDS-PAGE and Western blotting analysis. Data in A-D are representative of three independent experiments.

## Notes

### Competing Interest Statement

The authors have declared no competing interest.

